# Age-associated different transcriptome profiling in zebrafish and rat: insight into diversity of vertebrate aging

**DOI:** 10.1101/478438

**Authors:** Yusuke Kijima, Wang Wantong, Yoji Igarashi, Kazutoshi Yoshitake, Shuichi Asakawa, Yutaka Suzuki, Shugo Watabe, Shigeharu Kinoshita

## Abstract

**Background:** Aging and death are inevitable for most species and are of intense interest for human beings. Most mammals, including humans, show obvious aging phenotypes, for example, loss of tissue plasticity and sarcopenia. In this regard, fish provide attractive models because of their unique aging characteristics. First, the lifespan of fish is highly varied and some long-lived fish can live for over 200 years. Second, some fish show anti-aging features and indeterminate growth throughout their life. Because these characteristics are not found in mammalian model organisms, exploring mechanisms of senescence in fish is expected to provide new insights into vertebrate aging. Therefore, we conducted transcriptome analysis for brain, gill, heart, liver and muscle from 2-month-, 7-month-, 16month- and 39-month-old zebrafish. In addition, we downloaded RNA-seq data for sequential age related gene expression in brain, heart, liver and muscle of rat (1). These RNA-seq data from two species were compared, and common and species-specific features of senescence were analyzed.

**Results:** Screening of differentially expressed genes (DEGs) in all zebrafish tissues examined revealed up-regulation of circadian genes and down-regulation of *hmgb3a*. Comparative analysis of DEG profiles associated with aging between zebrafish and rat showed both conserved and clearly different aging phenomena. Furthermore, up-regulation of circadian genes with aging and down-regulation of collagen genes were observed in both species. On the other hand, in zebrafish, up-regulation of autophagy related genes in muscle and *atf3* in various tissues suggested fish-specific anti-aging characteristics. Consistent with our knowledge of mammalian aging, a tissue deterioration-related DEG profile was observed in rat. We also detected aging-associated down-regulation of muscle development and ATP metabolism-related genes in zebrafish gill. Correspondingly, hypoxia-related genes were systemically up-regulated in aged zebrafish, suggesting age-related hypoxia as a senescence modulator in fish.

**Conclusions:** Our results indicate both common and different aging profiles between fish and mammals. Gene expression profiles specific to fish will provide new insight for future translational research.

## Introduction

Senescence and death are inevitable for most species, but their features are diverse. Genes that contribute to aging phenomena have been widely screened by using established model organisms such as nematode, fly, and mouse. These studies have revealed conserved senescence-associated mechanisms involving Reactive Oxygen Species (ROS) and mechanistic Target of Rapamycin (mTOR) (2), and have identified various longevity genes, for example, sirtuin and klotho (3, 4). However, most of this research has been based on a small number of canonical model organisms; therefore, the diversity of senescence and aging that has evolved across species has been neglected. Exploring various species with differing aging phenotypes may, therefore, be of great value for better understanding of senescence. The recent advances in high-throughput sequencing technology make it possible to perform genome-wide gene screening of non-model organisms that show unique aging and senescence phenotypes. Indeed, several studies using noncanonical model organisms have suggested that various unique mechanisms are involved in the diversification of senescence (5-7).

In this regard, fish species are attractive models to study vertebrate senescence because of their unique aging characteristics. In comparison to mammals, fish show various anti-aging phenotypes. Mammals show apparent senescence and a determined lifespan. Progressive loss of plasticity of various tissues is a common feature in mammalian aging. In fish, however, various tissues retain high plasticity even in the adult stage. Adult fish can regenerate heart muscle (8, 9), whereas mammals lose this capacity soon after birth (10). Fish can increase skeletal muscle mass throughout their lifespan by both hypertrophic (size increase of existing muscle fibers) and hyperplastic (adding new muscle fibers) processes (11), whereas postnatal muscle growth in mammals occurs exclusively by hypertrophy (12). Fish have the longest known lifespans among vertebrates. Life spans exceeding 100 years were observed in several rockfish species by a combination of radio-isotope and growth ring measurements (13). The maximum age recorded for a rough-eye rockfish, *Sebastes aleutianus*, is 205 years (14). Furthermore, age estimation using radiocarbon dating of eye lens nuclei from the Greenland shark, *Somniosus microcephalus*, revealed its lifespan to be at least 272 years (15). Conversely, there are naturally short-lived fish, known as annual fish, such as sweetfish (*Plecoglossus altivelis*) and killifish which live less than 1 year. The turquoise killifish, *Nothobranchius furzeri*, has a 4-6 month lifespan and is currently the shortest-lived vertebrate that can be bred in captivity (16, 17). These unique features make fish an attractive model for understanding the diversity of vertebrate senescence and lifespan.

Zebrafish, *Danio rerio*, have been extensively studied as a vertebrate model organism because of easy rearing, short generation time, and transparency of embryos. In common with other fish species, zebrafish also have a highly regenerative capacity, even at the adult stage. Zebrafish can regrow injured tissues, such as fins (18), maxillary barbell (19), retinae (20), optic nerves (21), spinal cord (22), heart (23), brain (24), hair cells (25), pancreas (26), liver (27), and kidney (28). Dedifferentiation of mature myocytes is observed during regeneration of zebrafish extraocular muscle (29), whereas dedifferentiation of somatic cells is a very unusual phenomenon in mammals. Hyperplastic muscle growth continues in zebrafish at aged stages, as in other fish (30). The average lifespan of zebrafish is 3.5 years and aged zebrafish exhibit some deterioration of tissues, such as curvature of the spine, elevation of senescence associated β-galactosidase activity, increase of oxidized protein (31, 32), and decline in cognitive function (33-35), which are aging phenomena commonly found in mammals. The molecular bases of the above-mentioned anti-aging and aging characteristics are still ambiguous in zebrafish.

Here we present transcriptome analysis associated with systemic aging in zebrafish. We performed RNA-seq from brain, heart, liver, muscle and gill, at various growth stages. In addition, we conducted comparative analysis of our zebrafish data with age-associated RNA-seq data of rat. We discuss aging characteristics of fish in comparison with mammals.

## Results

### Overview of sequencing results

The quality filtering, mapping and expression quantification results of the sequencing data are summarized in Table 1. Approximately six million and 40 million reads per replicate were obtained from zebrafish and rat, respectively. Eighty-two percent of zebrafish reads and 96% of rat reads were mapped to their respective reference sequences. Assembly by Cufflinks detected approximately 40,000 genes in zebrafish and approximately 75,000 in rats. Of these genes, 26,381 in zebrafish and 29,552 in rat were annotated as known genes. Clustering using the expression levels of all the genes of each replicate showed that the sequence data was clustered by tissue in both zebrafish and rat (Additional file 1: Figure S1a, b). The skeletal muscle cluster was close to that of heart in both species. This is consistent with the fact that skeletal muscle and the heart are closely related tissues.

**Table 1:** Sequencing summary

|  | **Zebrafish** | **Rat** |
| --- | --- | --- |
| Type | Paired-end | Single-end |
| Sequence length | 100bp | 50bp |
| Reads per sample | 6019960 ($\pm 2071826$) | 42746365 ($\pm 10024371$) |
| Tissues | brain/heart/liver/muscle/gill | brain/heart/liver/muscle |
| Reference | GRCz10 | Rnor_6.0 |
| Mapping rate | 82.1% ($\pm 3.8$) | 95.5% ($\pm 1.2$) |
| Assembled genes | 42062 | 75910 |
| Annotated genes | 26381 | 29552 |
| Orthologous genes | 12,795 |  |

Values of reads per sample and mapping rate represented as the mean ± SD.

### Age-dependent DEGs among tissues in zebrafish

In each zebrafish tissue, comparison of gene expression between two growth stages was performed, and differentially expressed genes with a q-value ≦ 0.05 were defined as DEGs. Two genes, *hmgb3a* (High mobility group box 3a) and LOC566587 (ERBB receptor feedback inhibitor 1-like), were detected as DEGs in all five tissues (skeletal muscle, brain, heart, liver, and gill). To compare with rat data, we focused on the four common tissues, heart, skeletal muscle, brain and liver between rat and zebrafish. Figure 1 shows the distribution of DEGs among the four zebrafish tissues. Seventeen DEGs were commonly detected in the four tissues (Figure 1 and Additional file 2: Table S1). These 17 DEGs contain three circadian rhythm-related genes, *nr1d1*, *nr1d2a* and *sik1* (si: ch 211-235e 18.3 synonym) (36, 37).

**Figure 1.**
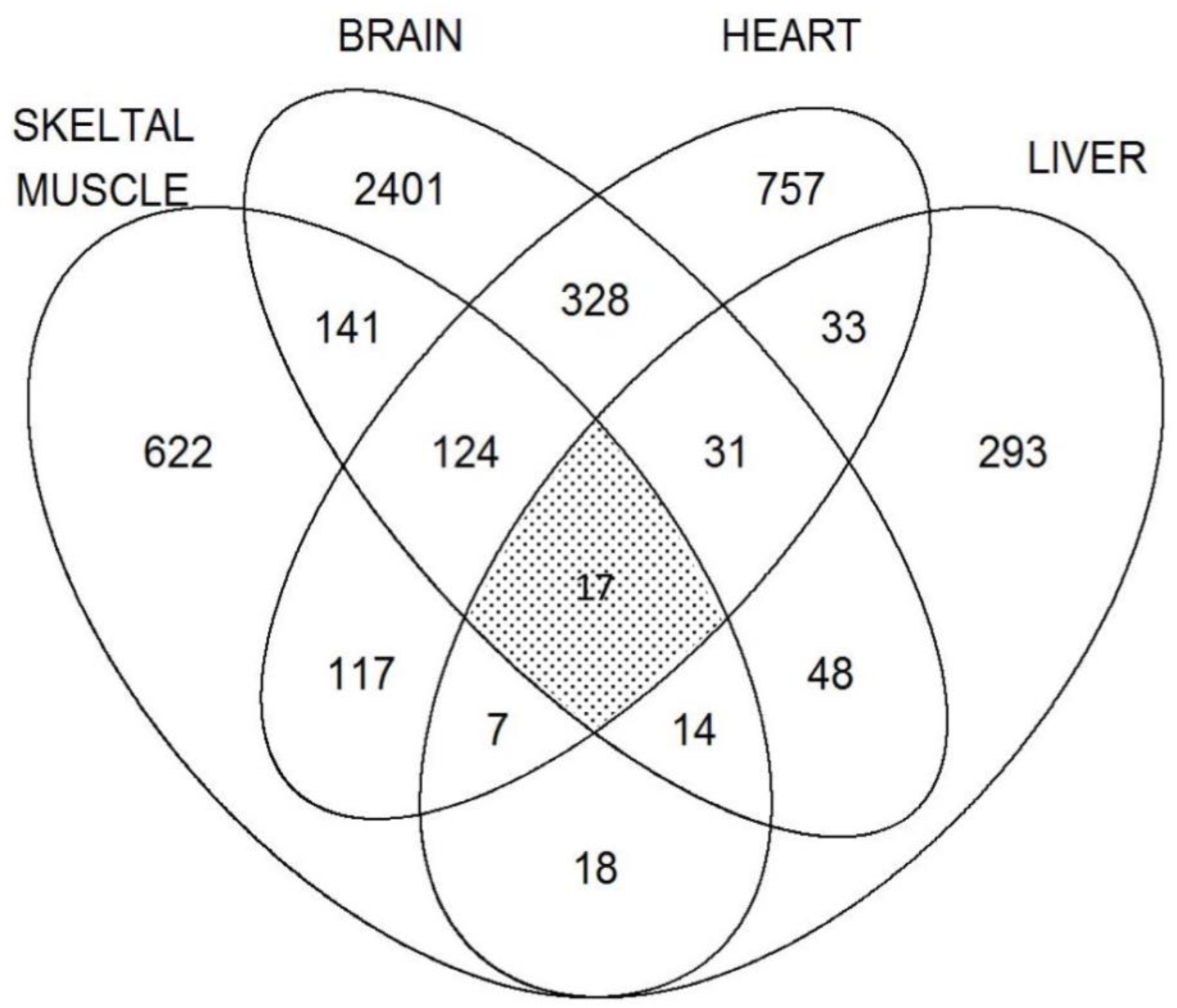
Venn diagram of zebrafish DEGs across four tissues. In each tissue, DEGs were detected by comparison between at least one pair from four growth stages (six pairs). 17 DEGs were commonly detected in all four tissues (meshed region).

To assess the age-dependent expression pattern of the 17 commonly detected DEGs, correlation coefficient matrixes were drawn for each tissue (Figure 2a - d). Most of the 17 genes showed similar expression changes with aging; only *hmgb3a* tended to inversely correlate in all tissues. Figures 2e and 2f show actual expression levels of these genes, with *hmgb3a* and the 16 other genes plotted separately. As shown in the figure, *hmgb3a* expression tended to decrease with aging whereas expression of the other 16 genes tended to increase. This age-associated decrease of *hmgb3a* expression was also observed in the gill (Additional file 2: Table S1).

**Figure 2.**
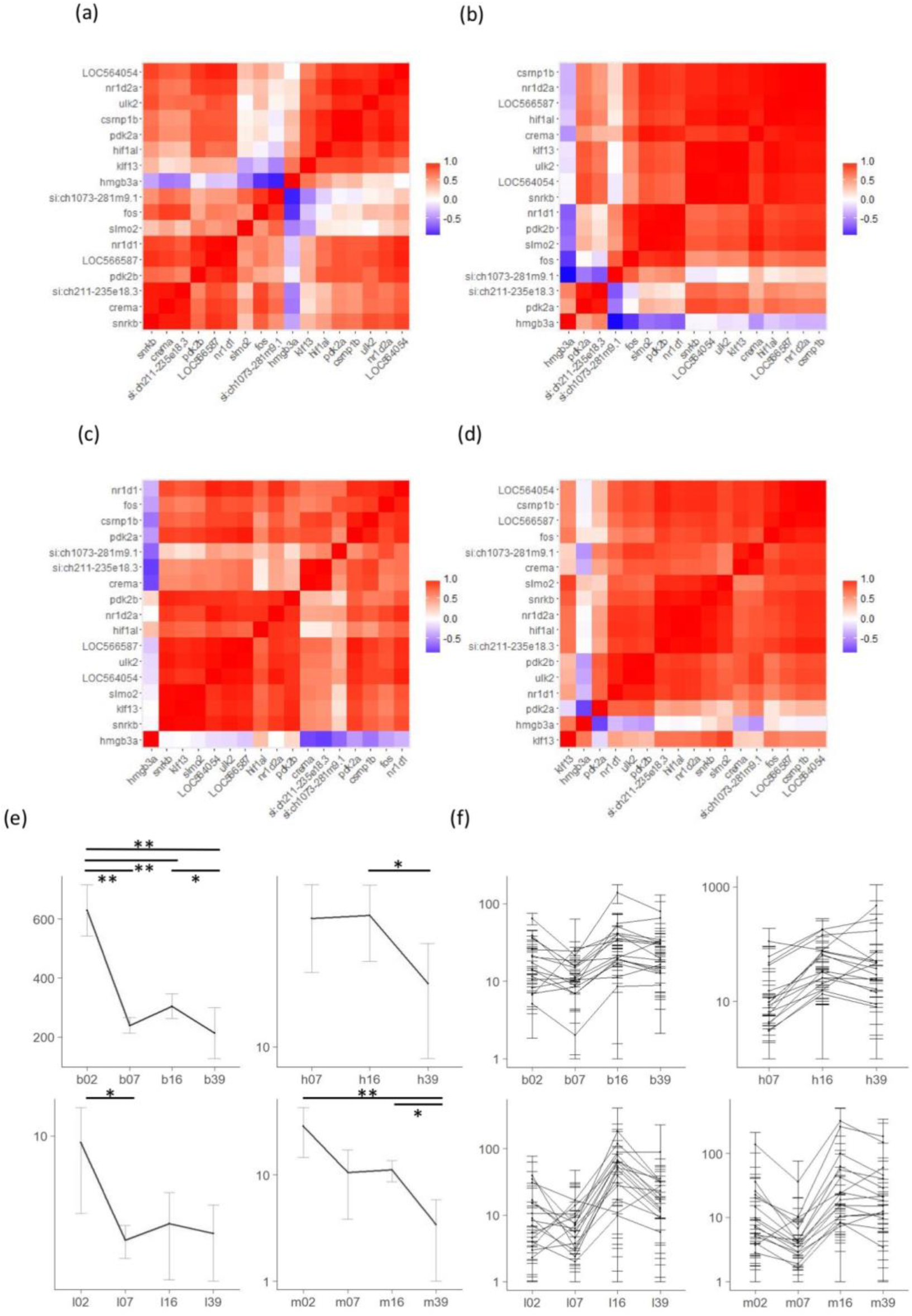
Expression patterns of DEGs common to zebrafish tissues. Correlation matrix of expression of 17 DEGs in brain (a), heart (b), liver (c), and muscle (d). Each cell indicates the correlation coefficient of growth/aging-dependent expression between each combination of the 17 DEGs commonly expressed in the four tissues. Most genes showed a correlated expression pattern, but *hmgb3a* showed an apparently different expression pattern in all four tissues. (f) Expression pattern of *hmgb3a*. In all four tissues, *hmgb3a* was expressed with the same decreasing level. (g) Expression pattern of DEGs expressed in all tissues, except *hmgb3a*. In all tissues, they showed a similar pattern, with a tendency of increasing expression. In all plots from (f) and (g), horizontal axes show growth stages. Labels “b”, “g”, “h”, “l”, “m” indicate “brain”, “gill”, “heart”, “liver” and “muscle”, respectively, and labels “02”, “07”, “16” and “39” indicate 2-, 7-, 16-, and 39-month-old zebrafish, respectively. For example, the label of “b02” means the brain of 2-month-old individuals. Vertical logarithmic axes show expression levels (FPKM) and error bars express a 95% confidence interval. **: q-value < 0.01; *: q-value < 0.05.

### Construction of orthologous gene set between zebrafish and rat

Based on the cross blastp screening between zebrafish and rat RNA-seq data, 12,795 genes were identified as orthologs (Table 1) and were designated as the Orthologous Gene Set (OGS). We used the OGS in the following comparative transcriptome analyses.

### Comparison of age-related gene expression patterns between zebrafish and rat using the OGS

To investigate time-course gene expression changes with aging, we focused on gene expression profiles at sequential growth stages of 7 months, 16 months and 39 months for zebrafish and 6 weeks, 21 weeks and 104 weeks for rat. To separate the OGSs by their expression patterns, we defined “D” as a significant decrease of gene expression, “U” as a significant increase and “M” as a change of −1 <log_2_ (Fold Change) <1. Then expression of each OGS from young to aged animals was divided into nine patterns, DD, DM, DU, MD, MM, MU, UD, UM and UU (Additional file 3: Figure S2). Figure 3a shows clustering based on the four patterns, DD, DU, UD, and UU. In the DD pattern (continuous decrease with age), some genes were clustered across muscle, heart and liver of rat (Figure 3a, cluster1). This cluster consisted of six genes, *Col6a3*, *Col4a5*, *Col1a1*, *Col1a2*, *Col5a1* and *Nid1*, all of which are collagen genes or a collagen-associated gene. Sequential expression patterns of these six genes were plotted in both species (Figure 3b, c). As in rat, zebrafish also showed a tendency for expression of collagen and collagen-related genes to decrease, but their expression was maintained up to the middle-age stage (Figure 3c), indicating robustness of zebrafish tissues against age-related down-regulation of collagen genes.

**Figure 3.**
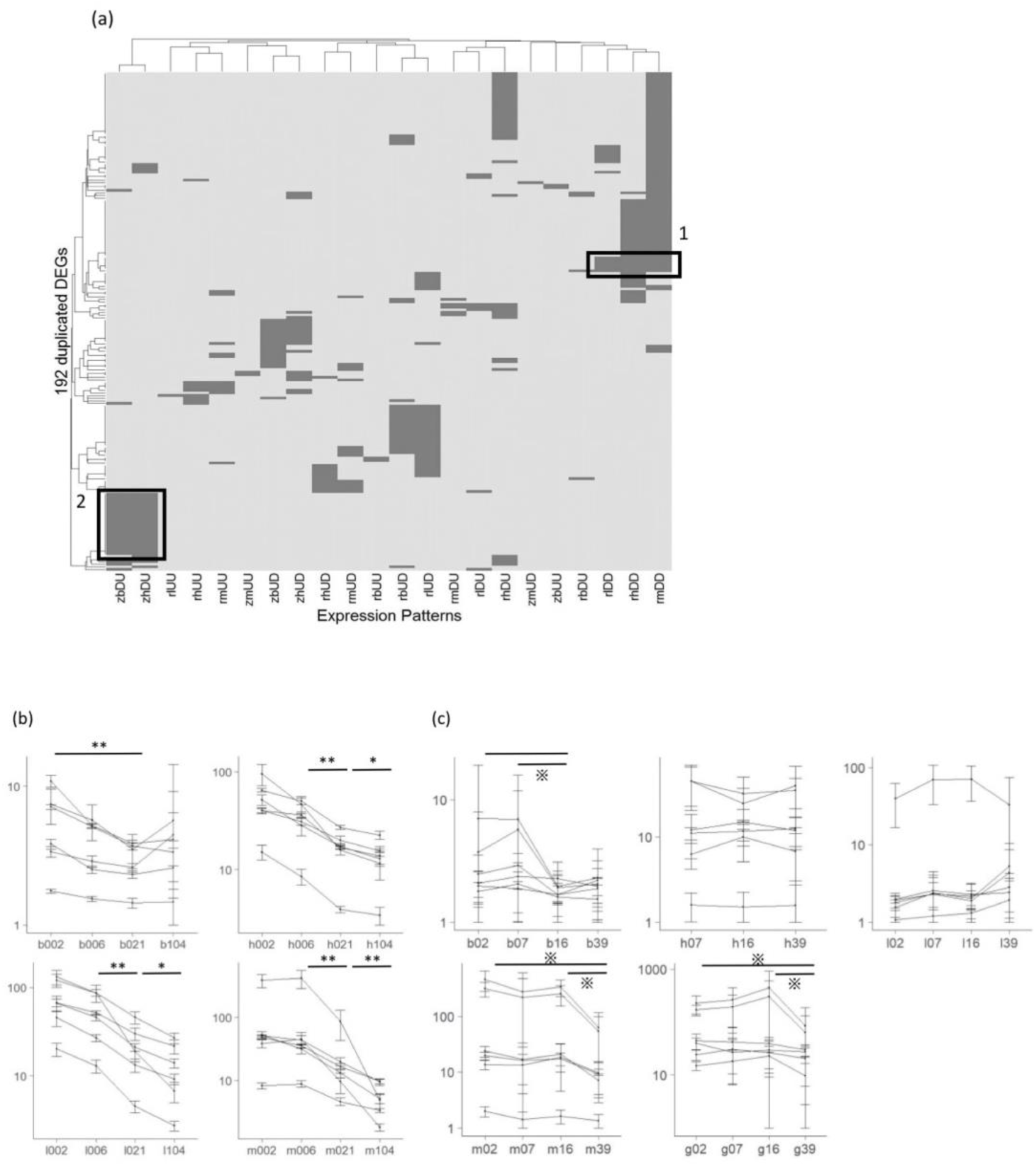
Analysis of sequential gene expression patterns of orthologous genes between rat and zebrafish. (a) Visualization of genes classified into four patterns (DD, DU, UD, UU) of each tissue for each species. For example, “zbDU” is a gene group that was down-regulated between 7 months and 16 months and up-regulated between 16 months and 39 months in the brain of zebrafish. “z” indicates zebrafish, r: rat, b: brain, h: heart, l: liver, m: muscle and g: gill. The horizontal axis is the age-related expression pattern for each tissue of a species and the vertical axis is genes duplicated in at least two groups. Colored cells show the corresponding gene is classified into the corresponding pattern. Genes commonly classified into “DD” in rat are in the black frame no.1 and those in “DU” in zebrafish brain and heart are in frame no.2. (b) The expression of six collagen genes, which are expressed in the four tissues, was down-regulated in rat and (c) zebrafish. Horizontal axes show growth stages. The zebrafish labels are the same as those in Figure 2 and the rat labels, “b”, “h”, “l” and “m” indicate “brain”, “heart”, “liver” and “muscle”, respectively, while “002”, “006”, “021” and “104” indicate 2-, 6-, 21- and 104-week-old rats, respectively. Vertical logarithmic axes show expression levels and error bars express 95% confidence intervals. **: q-value < 0.01 in ALL six genes; ※: q-value < 0.05 in ALL six genes; ※: q-value < 0.05 in SOME of six genes.

In the brain and heart of zebrafish, 25 genes formed a DU pattern cluster (up-regulated in middle-age then down-regulated in the aged stage) (Figure 3a, cluster 2). GO analysis with these 25 genes showed that hypoxia response and ribosome synthesis were significantly enriched (GO category: response to hypoxia; p=1.81E-002, GO category: ribosomal large subunit biogenesis; p=1.81E-002) (Table 2).

**Table 2.** GO analysis of DEGs regulated as “DU” in zebrafish

| Type | GO term | Adjusted p-value |
| --- | --- | --- |
| enriched | response to hypoxia | 1.81E-002 |
| enriched | response to oxygen levels&response to decreased oxygen levels | 1.81E-002 |
| enriched | ribosomal large subunit biogenesis | 1.81E-002 |
| enriched | nuclear transport&nucleocytoplasmic transport | 4.59E-002 |
| enriched | response to abiotic stimulus | 4.59E-002 |

GO analysis of 25 DEGs that were down-regulated between 7 months and 16 months and up-regulated between 16 months and 39 months was conducted. Genes related to response to hypoxia and ribosomal biogenesis were significantly enriched.

### Comparison of gene expression patterns between young and old stages in both species using the OGSs

To investigate senescence-related gene expression changes, OGSs were compared between young (7 months for zebrafish and 21 weeks for rat) and aged (39 months for zebrafish and 104 weeks for rat) stages. Based on previously reported age correspondence (38), 7- and 39-month-old zebrafish are equivalent to approximately 15- and 80-year-old humans, respectively, and 21- and 104-week-old rats are to approximately 15-20- and 60-year-old humans, respectively. All up-regulated and down-regulated DEGs were extracted from each tissue and each species. Then we detected 45 common DEGs between the two species and/or among two or more tissues. Figure 4 shows the 45 duplicated DEGs based on their expression patterns. We found four major clusters, up-regulated genes in both species, up-regulated genes in rat, up-regulated genes in zebrafish, and down-regulated genes in rat. *Per1*, *Per2*, *Tef* and *Bhle41* were up-regulated in the heart, liver and muscle of rat and the brain of zebrafish. These genes are related to the regulation of circadian rhythm (36, 39). Age-related up-regulation of circadian rhythm-related genes was also observed in zebrafish (Figure 4). Therefore, such up-regulation may be a common phenomenon between the two species.

**Figure 4.**
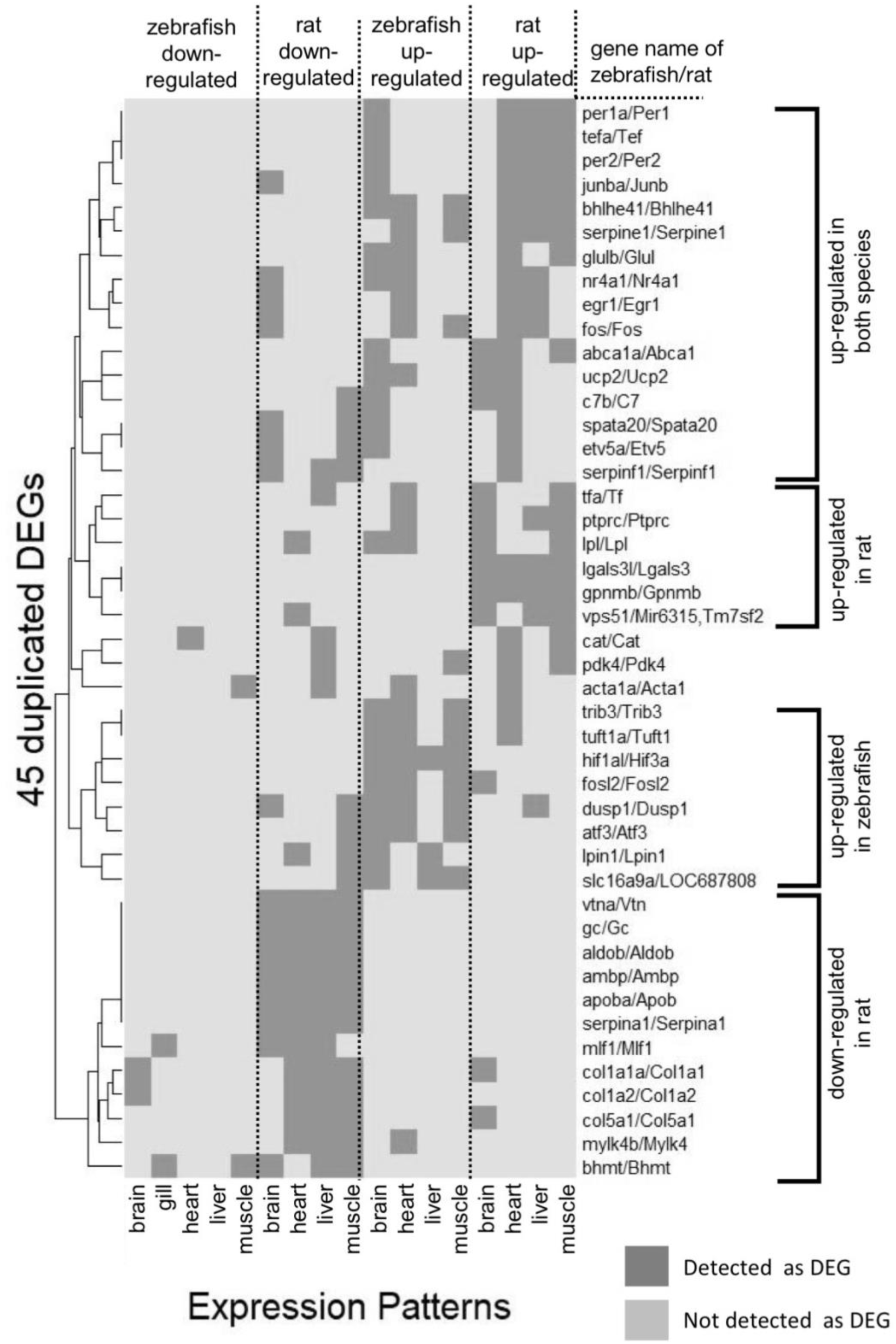
Analysis of DEGs in young vs old. Visualization of classified DEGs in young vs old. Horizontal axis is differential expression in each tissue in 21weeks vs 104 weeks in rat and 7 months vs 39 months in zebrafish. Vertical axis is genes duplicated at least once in the four groups.

Down-regulation of collagen genes, *Col1a1a*, *Col1a2*, and *Col5a1*, was observed in rat tissues (Figure 4). Such down-regulation of collagen genes were not obvious in zebrafish. These results are consistent with those in Figure 3.

Up-regulation of *fos*, *fosl2* and *atf3* were detected in multiple zebrafish tissues but not in rat (Figure 4). These genes are AP-1 transcription factors which are involved in various biological pathways. It is also known that these transcription factors are activated in response to hypoxia (40, 41). Genes responsive to hypoxia such as *trib3*, *hif1al* and *tuft1a* were up-regulated in various tissues in zebrafish (Figure 4), which is consistent with elevated expression of AP-1 transcription factors. It is noted that lifespan extension by inhibition of mTOR signaling in mouse caused up-regulation of ATF3 expression, which may indicate the remarkable feature of negligible senescence in fish (42).

**Figure 5.**
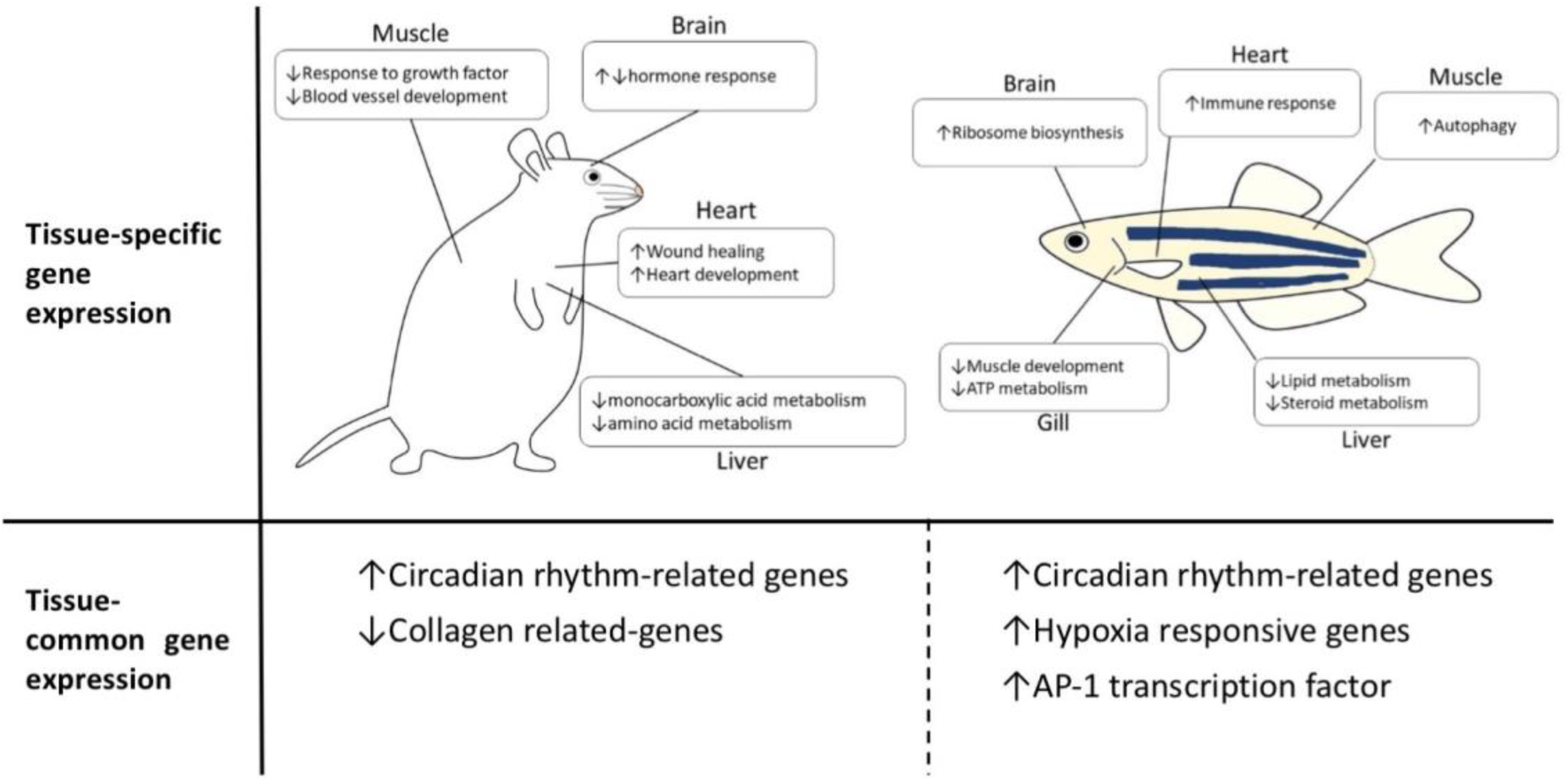
Summary of aging-associated gene expression profiles in zebrafish and rat. The upper row shows related biological process of tissue and species-specific DEGs and the lower row shows commonly expressed DEGs.

### GO analysis of tissue- and species-specific DEGs between young and old stages

To search for age-related DEGs that appear specifically in each tissue and species, DEG expression that was increased or decreased specifically in the brain, heart, liver, muscle or gill was extracted for both species. Age comparison was performed between young (7 months for zebrafish and 21 weeks for rat) and aged (39 months for zebrafish and 104 weeks for rat) stages. Subsequently, common DEGs between zebrafish and rat were excluded, and tissue- and species-specific DEGs were extracted from each comparison (Additional file 4: Figure S3). For the four tissues (excluding the gill), GO analysis of the tissue- and species-specific DEGs was performed (Additional files 5-8: Table S2-5) and is summarized in Table 3. As there is no rat data corresponding to the gill, GO analysis of the gill was conducted using gill-specific DEGs. Species-specific GO databases were used in the GO analysis.

Tissue-specific, up-regulated cascades in aged rat are listed in Additional file 5, Table S2. In brain, significantly up-regulated GO categories were ‘response to chemical substances’ such as hormones (GO category: response to peptide; p = 1.45 E-012, GO category: response to peptide steroid hormone; p = 6.36 E - 012) (Additional file 5: Table S2). In heart, ‘tissue repair’ and ‘development’ were up-regulated (GO category: wound healing; p = 1.16 E-008, GO category: heart development; p = 1.76 E-008). Expression of genes associated with ‘cell division’ was up-regulated in liver and muscle (GO category: cell division; p = 1.27 E - 016). ‘Lipid metabolism’, such as for monocarboxylic acid and fatty acids, was also significantly up-regulated in muscle (GO category: monocarboxylic acid metabolic process; p = 6.67 E - 006, GO category: fatty acid metabolic process; p = 1.20 E - 005).

Tissue-specific, down-regulated cascades in aged rat are listed in Additional file 6, Table S3. In brain, expression of genes associated with ‘hormone responses’ and ‘learning’ were significantly decreased (GO category: cellular response to hormone stimulus; p = 4.71 E-003, GO category: learning or memory; p = 1.37 E-002). In heart, ‘nucleotide metabolism’ was down-regulated (GO category: nucleotide metabolic process; p = 2.55 E - 017). In liver, ‘metabolism’ of amino acids and monocarboxylic acids, and ‘development’ (GO category: monocarboxylic acid metabolic process, p = 3.18E - 017, GO category: cellular amino acid metabolic process; p = 3.66E - 017, GO category: liver development; p = 4.02 E-009) were down-regulated. Also, in the muscle, expression of genes related to ‘blood vessel formation’ and ‘growth’ were down-regulated (GO category: blood vessel development; p = 2.90 E - 022, GO category: response to growth factor; p = 3.25 E - 017).

Tissue-specific, up-regulated cascades in zebrafish are listed in Additional file 7, Table S4. ‘Peptide biosynthesis’ and ‘ribosome synthesis’ were significantly up-regulated in brain (GO category: peptide biosynthetic process; p = 4.25 E-007, GO category: ribosomal large subunit assembly; p = 5.71 E- 007). Lifespan is prolonged by the reduction of ribosomal function; therefore, the up-regulation of the peptide and ribosome biosynthesis related genes is considered to accelerate aging (43, 44). In heart, there were significantly enriched categories associated with ‘activated immunity’ (GO category: positive regulation of immune system process; p = 1.25E - 002).

Tissue-specific, down-regulated cascades in zebrafish are listed in Additional file 8, Table S5. Genes down-regulated in liver were significantly enriched in categories related to ‘lipid metabolism’ and ‘steroid metabolism’ (GO category: lipid metabolic process; p = 7.44 E - 017, GO category: steroid metabolic process; p = 4.57 E-009). Notably, in gill, the expression of genes associated with ‘muscle development’ and ‘ATP metabolism’ were significantly down-regulated (GO category: muscle structure development; p = 5.05 E - 003, GO category: ATP metabolic process; p = 5.66 E - 003).

We also conducted GO analyses of zebrafish- and tissue-specific DEGs using the rat GO database (Additional files 9, 10: Table S6, 7). As a result, genes up-regulated in aged zebrafish muscles were significantly enriched in the category of ‘autophagy’ (GO category: autophagy; p = 6.24E-003). As in the analysis using the zebrafish GO database (Additional file 8: Table S5), down-regulation of ‘muscle contraction’ and ‘muscle formation’ related cascades in aged zebrafish was also observed in gill (Additional file 10: Table S7).

**Table 3.**
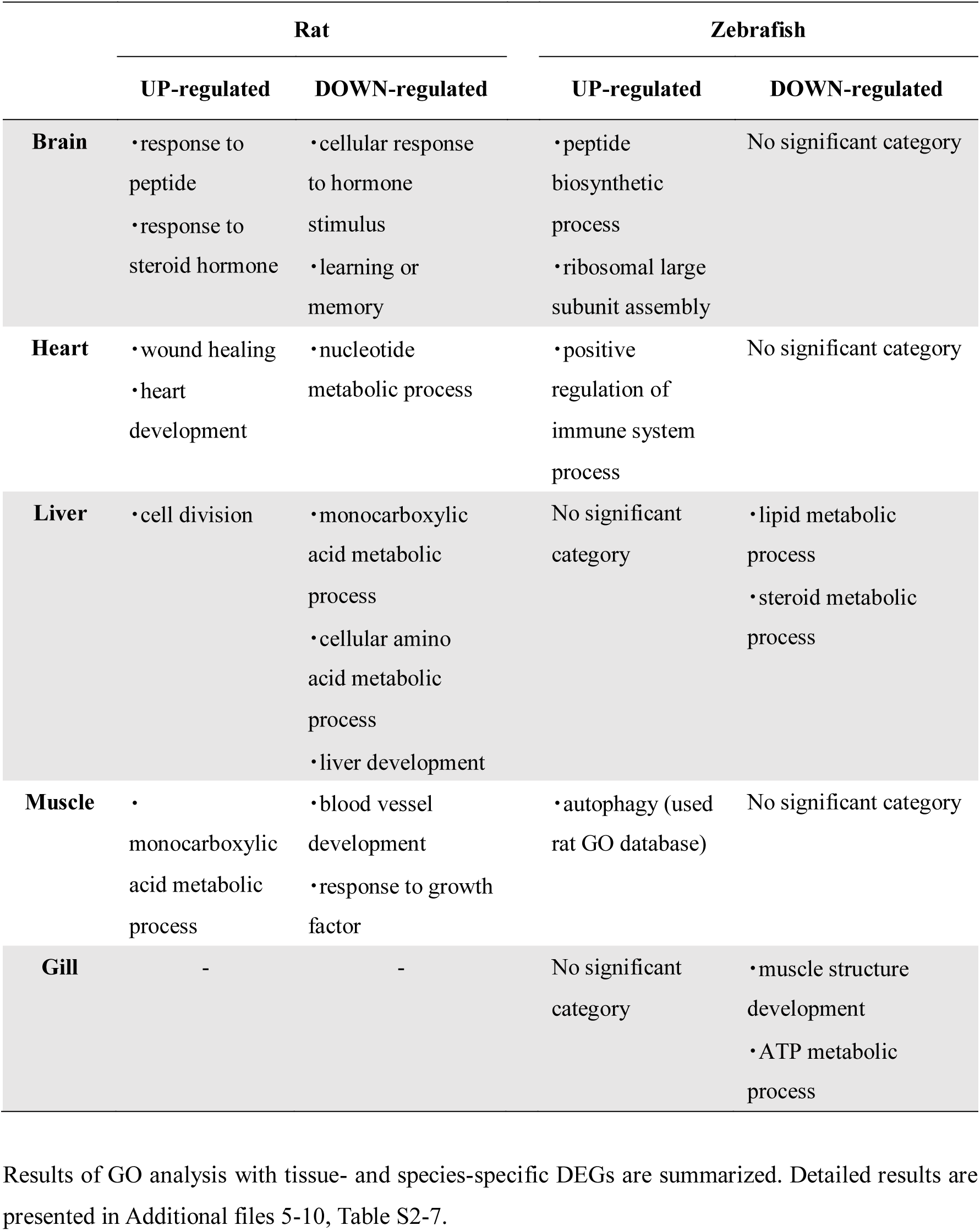
Summary of GO analysis with tissue and species specific DEGs

|  | Rat |  | Zebrafish |  |
| --- | --- | --- | --- | --- |
|  | UP-regulated | DOWN-regulated | UP-regulated | DOWN-regulated |
| **Brain** | • response to peptide<br>• response to steroid hormone | • cellular response to hormone stimulus<br>• learning or memory | • peptide biosynthetic process<br>• ribosomal large subunit assembly | No significant category |
| **Heart** | • wound healing<br>• heart development | • nucleotide metabolic process | • positive regulation of immune system process | No significant category |
| **Liver** | • cell division | • monocarboxylic acid metabolic process<br>• cellular amino acid metabolic process<br>• liver development | No significant category | • lipid metabolic process<br>• steroid metabolic process |
| **Muscle** | • monocarboxylic acid metabolic process | • blood vessel development<br>• response to growth factor | • autophagy (used rat GO database) | No significant category |
| **Gill** | - | - | No significant category | • muscle structure development<br>• ATP metabolic process |

Results of GO analysis with tissue- and species-specific DEGs are summarized. Detailed results are presented in Additional files 5-10, Table S2-7.

## Discussion

Special senescence and lifespan features make fish an interesting model for studying vertebrate aging. In this study, we conducted deep RNA sequencing of aging zebrafish. Our systematic transcript profiling across four growth stages, including an aged stage, provides unpreceded data in fish. In addition, we revealed common and different features of gene expression profiles with aging between fish and mammals by comparative transcriptome analysis using OGSs.

### Common features of aging-related changes to gene expression between zebrafish and rat

Up-regulation of circadian rhythm-related genes was commonly observed in zebrafish and rats. Many physiological processes follow circadian rhythms and the robustness of circadian rhythms deteriorates with age, including in zebrafish (32, 45). Solanas et al. (46) reported that the expression of genes responsible for the core circadian rhythm does not fluctuate with aging, but the components of gene set which follow circadian rhythm of aged mice do change. In addition, it has been shown that genes responsible for protein acetylation no longer show a circadian pattern of expression in the liver of aged mice (47). These reports imply that changes to gene expression in response to senescence are intimately related to circadian rhythm. Taken together, our results indicate a common age-related feature in both species; changes in the expression of circadian-related genes at the aged stage causes deterioration of the robustness of the circadian rhythm.

Age-associated repression of collagen genes in various tissues was also observed in both zebrafish and rat. Although collagen fibers accumulate in aged tissues of mammals, mRNA levels of collagen genes tend to decrease with aging (48, 49). Recent meta-analysis of age-related gene expression in mammals showed that reduction of collagen gene expression is a general feature of their aging (50). This suggests that age-related accumulation of collagen suppresses its gene expression to reduce the level of accumulation (49, 51). Although collagen accumulation in aged fish tissues has not been examined, our results indicate that age-associated reduction of collagen gene expression is a common feature of mammals and fish. However, reduction of collagen gene expression in zebrafish is more gradual than that in rat as shown in Figure 2c, where significant reduction was only observed in aged individuals.

### mTOR signaling

Mechanistic target of rapamycin (mTOR) signaling is a key modulator of growth and aging, and is conserved from yeast to human (52). Inhibition of mTOR signaling causes inhibition of senescence and lifespan extension in various organisms. Li et al.(42) reported slow-aging mice with elevated *atf3* and *atf4* expression by inhibition of mTOR signaling. Conversely, loss of *atf3* caused activation of mTOR signaling and its downstream S6K phosphorylation in mouse liver (53). In our RNA-seq analysis, *atf3* up-regulation was observed in aged zebrafish tissues, whereas no such up-regulation was observed in aged rat (see Figure 3). Moreover, aged rat muscle showed decreased expression of *atf3*. One of the downstream outputs of mTOR signaling is protein synthesis. ATF4 is a transcription factor that senses a deficit in protein synthesis and activates target genes including *atf3*. The *atf3* induction detected in this study indicates decreased mTOR signaling in aged zebrafish tissues. Resveratrol is a natural polyphenol that has various benefits for age-related mammalian diseases, such as diabetes, cancer, and neurodegenerative and cardiovascular diseases (54, 55). Hsu et al. (55) reported that resveratrol activates the anti-aging gene *klotho* via ATF3 activity.

Another downstream output of mTOR signaling is autophagy. Autophagy is negatively regulated by mTOR signaling (56). Consistent with *atf3* expression, which is also negatively regulated by mTOR signaling, we detected increased autophagy activity in aged zebrafish muscle (Table 3). Recent studies have revealed a close relationship between age-dependent decline of autophagy and senescence (56). Transgenic mice in which the autophagy inhibitory complex was disrupted show promoted longevity (57). Age-related deterioration of autophagy in muscle causes decreased regenerative capacity in mammalian muscle stem cells and it can be recovered by activation of autophagy (58). Mammalian skeletal muscles undergo marked senescence called sarcopenia, the loss of muscle mass and strength. Progressive loss of regenerative capacity of muscle is also a general feature of mammalian aging. Such a tendency is also detected in rat muscle in our study (Table 3). The mortality rate and pathogenesis of many age-related human diseases is associated with sarcopenia and the functional status of skeletal muscle (59, 60), suggesting that muscle is a key regulator of systemic aging. In fish, however, hyperplastic muscle growth (neo-muscle fiber formation) continues throughout life (11). The regeneration capacity of muscle is also high in zebrafish and adults can regenerate the heart after ventricle resection (23). The increased autophagy observed in this study may partly explain such anti-aging characteristics of fish muscle.

### Systemic decrease of *hmgb3a* expression in adult and aged zebrafish

A systemic decrease of *hmgb3a* expression was found to be a specific feature of zebrafish in comparison to rat. In all analyzed zebrafish tissues, *hmgb3a* expression gradually decreased with aging but was still detected in aged individuals (Figure 2f). *hmgb3a* is an orthologue of mammalian *Hmgb3* and belongs to the hmgb (High-Mobility Group Box) family. In mammals, *Hmgb3* shows high expression in the early developmental stage (61), but its expression is restricted into hematopoietic stem cells (HSCs) after birth. In zebrafish, however, systemic *hmgb3a* expression was detected even in the adult stage, in clear contrast to mammals. HMGB3 deficiency in mouse causes changes in the differentiation rate of the lymphoid and myeloid cells from HSCs (62), indicating HMGB3 function in the proper differentiation of HSCs. It is noted that aged mammalian HSCs show myeloid bias, a disrupted differentiation rate of lymphoid and myeloid cells from HSCs (63). Systemic expression of *hmgb3a* indicates that zebrafish *hmgb3a* functions in cells other than HSCs. Recent studies have revealed that hmgb family proteins in yeast and human are involved in mTOR signaling as general regulators of cell growth by controlling ribosome biogenesis (64).

### Age-associated hypoxia in zebrafish

Increased expression of hypoxia-responsive genes in aged individuals was observed in zebrafish tissues. Expression of AP-1 transcription factors was also up-regulated. These results suggest that the aged zebrafish is in a hypoxic state, which is consistent with the predicted deterioration of gill function from gills-specific down-regulation of muscle development and ATP metabolism-related genes (Table 3). The muscles present in the gill are the abductor muscle and the adductor muscle and fish efficiently take up oxygen by combining the movements of these muscles (65). Here, we suggest that gill-specific down-regulation of genes related to muscle development is associated with deterioration of gill function of zebrafish, which leads to systemic hypoxia.

Hypoxia and ischemia induce mitochondrial production of reactive oxygen species (ROS). It has long been proposed that ROS accelerate aging by inflicting damage on molecules such as proteins, lipids, and DNA. In rats, age-related expansion of hypoxia in the kidney has been reported. The degree of hypoxia in the kidney correlated with age-related tubulointerstitial injury (66).

Despite various anti-aging characteristics, the lifespan of zebrafish is 3-5 years. Age-associated decline of gill function and resulting hypoxia may be a trigger of senescence in zebrafish. Chronic hypoxia effects adult fish physiology and can cause pathological conditions. For example, exposure of adult zebrafish to hypoxia for 11 days caused retinopathy, an angiogenesis-dependent disease (67). However, effects of hypoxia on aging and lifespan in fish have not been examined.

## Conclusions

The age-related gene expression profiles in zebrafish and rat are summarized in Figure 4. Our analysis revealed similarities and differences in age-related gene expression profiles between zebrafish and rat. Both species showed age-associated changes in expression of genes related to circadian rhythm. However, increased expression of *atf3* and up-regulation of autophagy in aged zebrafish was in clear contrast to the situation in rat. These changes suggest down-regulation of mTOR signaling in aged zebrafish. Taken together with the systemic expression of *hmgb3a*, these features may explain anti-aging characteristics observed in fish. Notably, the expression of AP-1 transcription factor and hypoxia responsive genes are elevated in multiple tissues of aged zebrafish. Although it is necessary to examine whether hypoxia affects the aging of zebrafish, age-related hypoxia may be a senescence modulator in fish. This is the first report presenting deep RNA-seq of various tissues in aging fish. Fish consist of more than 30,000 species, the most diverse vertebrate group. Their lifespan also varies extremely depending on the species, ranging from over hundreds of years to a few months. Our zebrafish data and future comparative analyses with long-lived or short-lived fish species will provide new insight into the diversity of vertebrate aging and lifespan.

## Methods

### Tissue collection from zebrafish

Brain, heart, liver, muscle and gill tissues were collected from zebrafish at four growth stages of 2, 7, 16 and 39 months of age. Four to five replicates were prepared per experimental group. The heart was not collected at 2 months because of technical difficulty. All zebrafish were bred in tanks with circulated water at 28.5°C with a lighting cycle of 9:30 on and 23:00 off. The sampling time was 14:00-15:00 for 2 months, 15:30~16:30 for 7 months, 13:00~14:00 for 16 months and 11:00~12:00 for 39 months.

### RNA extraction and construction of cDNA libraries

Total RNA was extracted from each sample using an RNeasy Mini Kit (QIAGEN), and cDNA libraries were constructed using a TruSeq Stranded mRNA HT Kit from Illumina from a total of 94 samples. cDNA concentration was determined by qPCR using the KAPA SYBR^®^ FAST qPCR system (library specific primers: 5’- AAT GAT ACG GCG ACC GA −3’, 5’- CAA GCA GAA GAC GC ATA CGA −3’).

### Sequencing and subsequent processes

cDNA libraries were subjected to paired-end sequencing with the Illumina Hiseq 2000 system. Poly A tails were removed from each read using PRINSEQ-lite 0.20.4 and quality filtering was performed using FASTX Toolkit 0.0.13 (http://hannonlab.cshl.edu/fastx_toolkit/) (68). These filtered reads were mapped onto the zebrafish reference genome GRCz10 using Tophat2 version 2.1.1 and assembled with Cufflinks 2.2.1 (69, 70). We also used Cuffdiff in the Cufflinks package to obtain differentially expressed genes (DEGs) between groups from the four growth stages with q-value <0.05 (71). After that, detailed analysis of data was performed using cummeRbund of R package (72). For Gene ontology (GO) analysis, we used GeneTrail 2 version 1.5 (https://genetrail2.bioinf.uni-sb.de/start.html) (73).

### Obtaining rat RNA-seq data

Rat RNA-seq data set for the brain, heart, liver and muscle (Accession Number: SRP037986) was downloaded from DDBJ (http://www.ddbj.nig.ac.jp/searches-j.html) in fastq format. Subsequent analysis was conducted with the same pipeline used for zebrafish, as described above. Reference files of both zebrafish and rat used for mapping and assembly were downloaded from iGenomes (https://support.illumina.com/sequencing/sequencing_software/genome.html).

### Extraction of orthologous genes between rat and zebrafish

To detect orthologous genes between rat and zebrafish, the sequences of all transcripts of both species derived from cufflinks were translated into amino acid sequences in six reading frames with transeq in EMBOSS version 6.6.0.0 (74, 75). Subsequently, homology searches between rat and zebrafish were performed on the amino acid sequences using blastp in BLAST+ 2.6.0 (76). From this BLAST result, only transcripts having homology of e-value <1.0e-30 were retained and these homology searches were performed bi-directionally. The highest ranking transcripts from the homology searches in both directions were converted to genes and defined as orthologous genes. When one gene of one species hit two genes of the other species, one pair was defined as one orthologous gene.

### Declarations

#### Ethics approval and consent to participate

All our experiments were approved by the institutional animal ethics guidelines of the University of Tokyo.

#### Consent for publication

Not applicable

#### Acailability of data and materials

RNA-seq data of zebrafish generated and analysed during the current study are available in the DDBJ (https://www.ddbj.nig.ac.jp), accession [##Now submitting##].

The datasets of RNA-seq of rat are available in the DDBJ, accession SRP037986.

#### Author’s contributions

Study conception and design: YK, WW, SA, SW and SK. Sample collection: WW. RNA extraction: WW. Library construction and sequencing: YK, WW and YS. Analysis and interpretation of NGS data: YK, YI, KY. Drafting manuscript: YK. Critical revision: SK. All authors read and approved the final manuscript.

## Supporting information

## Acknowledgements

We thank Jeremy Allen, PhD, from Edanz Group (www.edanzediting.com/ac) for editing a draft of this manuscript.

## Additional files

### Additional file 1: Figure S1

∙ PDF
∙ Cluster analysis based on gene expression of all sample.
∙ Hierarchical clustering from gene expression in each replicate of zebrafish (a) and rat (b). “b” : brain, “h” : heart, “l” : liver, “m” : muscle and “g” : gill. All expression values were log2- transformed and each sample was clustered with Pearson’s correlation.

### Additional file 2: Table S1

∙ DOCX
∙ DEGs common to brain, heart, liver and muscle in zebrafish.
∙ DEGs common to 4 tissues of brain, heart, liver and muscle are summarized. Significantly upregulated genes are colored with red and downregulated are with blue in seventh column. Two genes down-regulated in gill were added to last. In third and fourth column, the compared experimental groups are subscribed. “b”: brain, “h”: heart, “l”: liver, “m”: muscle and “g”: gill and “02”: 2 months after birth, “07”: 7 months, “16”: 16 months and “39”: 39 months.

### Additional file 3: Figure S2

∙ JPG
∙ Summary of genes classified into 9 expression patterns.
∙ Expression of the orthologous genes between rat and zebrafish were classified into 9 patterns. “U” means the expression level increases between growth stages, “D” does decreases and “N” does the level does not significantly alter. The early growth stage (2 weeks of rat and 2 months of zebrafish) was excepted in this analysis. Blue and red colored cells contain relatively small and large numbers of genes, respectively.

### Additional file 4: Figure S3

∙ PDF
∙ Procedure to extract tissue- and species-specific DEGs between young and old stages.
∙ Left : Tissue-specific DEGs which were <u>down-regulated</u> in each tissue of zebrafish and rat. DEGs in zebrafish were compared to DEGs in the same tissue of rat. Right : Tissue-specific DEGs which were <u>up-regulated</u> in each tissue of zebrafish and rat. These DEGs in zebrafish were compared to DEGs in the same tissue of rat.

### Additional file 5: Table S2

∙ DOCX
∙ GO analysis with up-regulated genes in rat
∙ (a) brain, (b) heart, (c) liver, (d) muscle. 1st~15th significant categories were selected if the significantly enriched categories are so much. All analysis was conducted with GeneTrail2 version 1.5.

### Additional file 6: Table S3

∙ DOCX
∙ GO analysis with down-regulated genes in rat
∙ (a) brain, (b) heart, (c) liver, (d) muscle. 1st~15th significant categories were selected if the significantly enriched categories are so much. All analysis was conducted with GeneTrail2 version 1.5.

### Additional file 7: Table S4

∙ DOCX
∙ GO analysis with up-regulated genes in zebrafish
∙ (a) brain, (b) heart, (c) liver, (d) muscle, (e) gill. 1st~15th significant categories were selected if the significantly enriched categories are so much. All analysis was conducted with GeneTrail2 version 1.5.

### Additional file 8: Table S5

∙ DOCX
∙ GO analysis with down-regulated genes in zebrafish
∙ (a) brain, (b) heart, (c) liver, (d) muscle, (e) gill. 1st~15th significant categories were selected if the significantly enriched categories are so much. All analysis was conducted with GeneTrail2 version 1.5.

### Additional file 9: Table S6

∙ DOCX
∙ GO analysis of up-regulated genes in zebrafish which are converted into corresponding genes in rat
∙ Zebrafish DEGs of each tissue are converted into orthologous genes of rat. (a) brain, (b) heart, (c) liver, (d) muscle, (e) gill. 1st~15th significant categories were selected if the significantly enriched categories are so much. All analysis was conducted with GeneTrail2 version 1.5.

### Additional file 10: Table S7

∙ DOCX
∙ GO analysis with down-regulated genes in zebrafish which are converted into corresponding genes in rat
∙ Zebrafish DEGs of each tissue are converted into orthologous genes of rat. (a) brain, (b) heart, (c) liver, (d) muscle, (e) gill. 1st~15th significant categories were selected if the significantly enriched categories are so much. All analysis was conducted with GeneTrail2 version 1.5.

