## Supplementary material for "Age-associated different transcriptome profiling in zebrafish and rat: insight into diversity of vertebrate aging"

(a) zebrafish_brain_DOWN

| **Type** | **Name** | **Adjusted p-value** |
| --- | --- | --- |
| enriched | axon development | 1.49E-002 |
| enriched | cellular response to growth factor stimulus | 1.49E-002 |
| enriched | neuron projection morphogenesis | 1.49E-002 |
| enriched | response to growth factor | 1.49E-002 |
| enriched | positive regulation of canonical Wnt signaling pathway | 1.86E-002 |
| enriched | cell morphogenesis involved in neuron differentiation | 2.82E-002 |

(b) zebrafish_heart_DOWN

| **Type** | **Name** | **Adjusted p-value** |
| --- | --- | --- |
| enriched | tube development | 3.40E-005 |
| enriched | embryonic organ development | 1.29E-004 |
| enriched | tissue morphogenesis | 1.29E-004 |
| enriched | gland development | 2.25E-004 |
| enriched | muscle structure development | 4.55E-004 |
| enriched | vasculature development | 4.55E-004 |
| enriched | heart growth | 4.68E-004 |
| enriched | embryonic morphogenesis | 4.99E-004 |
| enriched | collagen fibril organization | 9.05E-004 |
| enriched | morphogenesis of an epithelium | 9.05E-004 |
| enriched | reproductive structure development | 9.05E-004 |
| enriched | reproductive system development | 9.05E-004 |
| enriched | muscle organ development | 1.02E-003 |
| enriched | urogenital system development | 1.02E-003 |
| enriched | cardiac chamber morphogenesis | 1.09E-003 |

(c) zebrafish_liver_DOWN

| **Type** | **Name** | **Adjusted p-value** |
| --- | --- | --- |
| enriched | sterol biosynthetic process | 2.91E-022 |
| enriched | secondary alcohol biosynthetic process & cholesterol biosynthetic process | 9.51E-020 |
| enriched | sterol metabolic process | 1.55E-018 |
| enriched | alcohol biosynthetic process | 6.49E-018 |
| enriched | steroid biosynthetic process | 7.09E-018 |
| enriched | alcohol metabolic process | 5.79E-017 |
| enriched | cholesterol metabolic process | 2.26E-016 |
| enriched | organic hydroxy compound biosynthetic process | 2.26E-016 |
| enriched | secondary alcohol metabolic process | 2.55E-016 |
| enriched | steroid metabolic process | 2.78E-015 |
| enriched | organic hydroxy compound metabolic process | 3.25E-015 |
| enriched | lipid biosynthetic process | 7.58E-015 |
| enriched | small molecule biosynthetic process | 6.16E-013 |
| enriched | isoprenoid biosynthetic process | 1.23E-006 |
| enriched | phospholipid metabolic process | 3.14E-006 |

(d) zebrafish_muscle_DOWN

| **Type** | **Name** | **Adjusted p-value** |
| --- | --- | --- |
| No significant categories | | |

(e) zebrafish_gill_DOWN

| **Type** | **Name** | **Adjusted p-value** |
| --- | --- | --- |
| enriched | striated muscle contraction | 1.33E-008 |
| enriched | muscle system process | 1.76E-008 |
| enriched | muscle contraction | 2.90E-008 |
| enriched | skeletal muscle contraction | 4.55E-007 |
| enriched | striated muscle tissue development | 5.81E-007 |
| enriched | muscle tissue development | 7.79E-007 |
| enriched | multicellular organismal movement & musculoskeletal movement | 9.70E-007 |
| enriched | muscle structure development | 1.49E-006 |
| enriched | regulation of muscle contraction | 6.31E-006 |
| enriched | striated muscle cell development | 6.51E-006 |
| enriched | muscle cell development | 9.96E-006 |
| enriched | regulation of muscle system process | 3.91E-005 |
| enriched | muscle organ development | 6.75E-005 |
| enriched | striated muscle cell differentiation | 1.89E-004 |
| enriched | regulation of striated muscle contraction | 3.43E-004 |
