## Supplementary material for "Age-associated different transcriptome profiling in zebrafish and rat: insight into diversity of vertebrate aging"

(a)zebrafish_brain_UP

| **Type** | **Name** | **Adjusted p-value** |
| --- | --- | --- |
| enriched | response to alcohol | 1.65E-004 |
| enriched | response to peptide hormone | 1.65E-004 |
| enriched | response to radiation | 1.65E-004 |
| enriched | ribosomal large subunit assembly | 1.65E-004 |
| enriched | response to light stimulus | 2.11E-004 |
| enriched | response to peptide | 3.29E-004 |
| enriched | response to steroid hormone | 3.52E-004 |
| enriched | response to drug | 3.77E-004 |
| enriched | response to extracellular stimulus | 4.96E-004 |
| enriched | ribosomal large subunit biogenesis | 5.15E-004 |
| enriched | ribosome biogenesis | 5.77E-004 |
| enriched | cellular response to hormone stimulus | 6.16E-004 |
| enriched | small molecule biosynthetic process | 6.16E-004 |
| enriched | response to corticosterone | 8.45E-004 |
| enriched | ribosome assembly | 8.96E-004 |

(b)zebrafish_heart_UP

| **Type** | **Name** | **Adjusted p-value** |
| --- | --- | --- |
| enriched | leukocyte activation | 8.60E-003 |
| enriched | lymphocyte activation | 1.63E-002 |
| enriched | positive regulation of cell proliferation | 1.94E-002 |
| enriched | positive regulation of immune system process | 1.94E-002 |
| enriched | regulation of leukocyte proliferation | 1.94E-002 |
| enriched | regulation of lymphocyte proliferation | 1.94E-002 |
| enriched | regulation of mononuclear cell proliferation | 1.94E-002 |
| enriched | respiratory burst | 1.94E-002 |
| enriched | cell activation involved in immune response | 4.12E-002 |
| enriched | leukocyte activation involved in immune response | 4.12E-002 |
| enriched | regulation of protein serine threonine kinase activity | 4.12E-002 |
| enriched | regulation of cytoplasmic transport | 4.56E-002 |
| enriched | cellular cation homeostasis | 4.94E-002 |
| enriched | cellular ion homeostasis | 4.94E-002 |
| enriched | immune effector process | 4.94E-002 |

(c)zebrafish_liver_UP

| **Type** | **Name** | **Adjusted p-value** |
| --- | --- | --- |
| No significant categories | | |

(d)zebrafish_muscle_UP

| **Type** | **Name** | **Adjusted p-value** |
| --- | --- | --- |
| enriched | autophagy | 6.24E-003 |

(e)zebrafish_gill_UP

| **Type** | **Name** | **Adjusted p-value** |
| --- | --- | --- |
| No significant categories | | |
