## Supplementary material for "Age-associated different transcriptome profiling in zebrafish and rat: insight into diversity of vertebrate aging"

(a) zebrafish_brain_DOWN

| **Type** | **Name** | **Adjusted p-value** |
| --- | --- | --- |
| No significant categories | | |

(b) zebrafish_heart_DOWN

| **Type** | **Name** | **Adjusted p-value** |
| --- | --- | --- |
| No significant categories | | |

(c) zebrafish_liver_DOWN

| **Type** | **Name** | **Adjusted p-value** |
| --- | --- | --- |
| enriched | lipid metabolic process | 7.44E-017 |
| enriched | lipid biosynthetic process | 1.84E-012 |
| enriched | cellular lipid metabolic process | 2.40E-010 |
| enriched | steroid metabolic process | 4.57E-009 |
| enriched | alcohol metabolic process | 8.84E-009 |
| enriched | steroid biosynthetic process | 4.27E-008 |
| enriched | sterol metabolic process | 4.27E-008 |
| enriched | organic hydroxy compound metabolic process | 4.54E-008 |
| enriched | sterol biosynthetic process | 7.26E-007 |
| enriched | alcohol biosynthetic process | 1.68E-006 |
| enriched | organic hydroxy compound biosynthetic process | 9.91E-006 |
| enriched | small molecule biosynthetic process | 2.68E-005 |
| enriched | isoprenoid biosynthetic process | 1.41E-004 |
| enriched | isoprenoid metabolic process | 1.13E-003 |
| enriched | organophosphate metabolic process | 5.30E-003 |

(d) zebrafish_muscle_DOWN

| **Type** | **Name** | **Adjusted p-value** |
| --- | --- | --- |
| No significant categories | | |

(e) zebrafish_gill_DOWN

| **Type** | **Name** | **Adjusted p-value** |
| --- | --- | --- |
| enriched | muscle structure development | 5.05E-003 |
| enriched | ATP metabolic process | 5.66E-003 |
| enriched | generation of precursor metabolites and energy | 5.66E-003 |
| enriched | nucleoside monophosphate metabolic process | 5.66E-003 |
| enriched | nucleoside triphosphate metabolic process | 5.66E-003 |
| enriched | purine nucleoside monophosphate metabolic process&purine ribonucleoside monophosphate metabolic process | 5.66E-003 |
| enriched | purine nucleoside triphosphate metabolic process | 5.66E-003 |
| enriched | purine ribonucleoside triphosphate metabolic process | 5.66E-003 |
| enriched | ribonucleoside monophosphate metabolic process | 5.66E-003 |
| enriched | ribonucleoside triphosphate metabolic process | 5.66E-003 |
| enriched | single organism carbohydrate catabolic process | 5.66E-003 |
| enriched | carbohydrate catabolic process | 5.83E-003 |
| enriched | purine nucleoside metabolic process | 9.09E-003 |
| enriched | purine ribonucleoside metabolic process | 9.09E-003 |
| enriched | ribonucleoside metabolic process | 1.29E-002 |
