## Supplementary material for "Age-associated different transcriptome profiling in zebrafish and rat: insight into diversity of vertebrate aging"

(a)zebrafish_brain_UP

| **Type** | **Name** | **Adjusted p-value** |
| --- | --- | --- |
| enriched | peptide biosynthetic process | 4.25E-007 |
| enriched | amide biosynthetic process | 4.59E-007 |
| enriched | cellular amide metabolic process | 4.59E-007 |
| enriched | peptide metabolic process | 4.59E-007 |
| enriched | ribosomal large subunit assembly | 5.71E-007 |
| enriched | translation | 6.25E-007 |
| enriched | ribosomal large subunit biogenesis | 1.51E-005 |
| enriched | ribosome assembly | 1.51E-005 |
| enriched | ribosome biogenesis | 3.27E-004 |
| enriched | ribonucleoprotein complex biogenesis | 3.75E-003 |
| enriched | ribonucleoprotein complex assembly | 4.30E-003 |
| enriched | ribonucleoprotein complex subunit organization | 6.23E-003 |

(b)zebrafish_heart_UP

| **Type** | **Name** | **Adjusted p-value** |
| --- | --- | --- |
| enriched | cell activation | 1.25E-002 |
| enriched | locomotion | 1.25E-002 |
| enriched | positive regulation of immune system process | 1.25E-002 |
| enriched | cell migration | 4.30E-002 |
| enriched | cell motility&localization of cell | 4.30E-002 |
| enriched | regulation of immune system process | 4.30E-002 |
| enriched | response to external stimulus | 4.30E-002 |

(c)zebrafish_liver_UP

| **Type** | **Name** | **Adjusted p-value** |
| --- | --- | --- |
| No significant categories | | |

(d)zebrafish_muscle_UP

| **Type** | **Name** | **Adjusted p-value** |
| --- | --- | --- |
| No significant categories | | |

(e)zebrafish_gill_UP

| **Type** | **Name** | **Adjusted p-value** |
| --- | --- | --- |
| No significant categories | | |
