## Supplementary material for "Age-associated different transcriptome profiling in zebrafish and rat: insight into diversity of vertebrate aging"

(a) rat_brain_DOWN

| **Type** | **Name** | **Adjusted p-value** |
| --- | --- | --- |
| enriched | cellular response to hormone stimulus | 4.71E-003 |
| enriched | response to peptide hormone | 4.71E-003 |
| enriched | response to steroid hormone | 4.71E-003 |
| enriched | response to corticosterone | 6.03E-003 |
| enriched | response to peptide | 6.03E-003 |
| enriched | learning or memory | 1.37E-002 |
| enriched | response to mineralocorticoid | 1.37E-002 |
| enriched | regulation of synapse structure or activity | 1.56E-002 |
| enriched | cognition | 2.05E-002 |
| enriched | skeletal muscle cell differentiation | 2.43E-002 |
| enriched | response to alcohol | 4.79E-002 |

(b) rat_heart_DOWN

| **Type** | **Name** | **Adjusted p-value** |
| --- | --- | --- |
| enriched | nucleobase containing small molecule metabolic process | 1.83E-017 |
| enriched | nucleotide metabolic process | 2.55E-017 |
| enriched | nucleoside phosphate metabolic process | 2.57E-017 |
| enriched | generation of precursor metabolites and energy | 7.47E-017 |
| enriched | cellular respiration | 1.79E-016 |
| enriched | energy derivation by oxidation of organic compounds | 1.79E-016 |
| enriched | small molecule catabolic process | 5.68E-015 |
| enriched | purine nucleoside metabolic process | 1.41E-014 |
| enriched | purine ribonucleoside metabolic process | 9.08E-014 |
| enriched | nucleoside metabolic process | 9.99E-014 |
| enriched | organic acid catabolic process&carboxylic acid catabolic process | 2.57E-013 |
| enriched | ribonucleoside metabolic process | 2.67E-013 |
| enriched | purine ribonucleotide metabolic process | 2.79E-013 |
| enriched | glycosyl compound metabolic process | 2.94E-013 |
| enriched | purine nucleotide metabolic process | 4.77E-013 |

(c) rat_liver_DOWN

| **Type** | **Name** | **Adjusted p-value** |
| --- | --- | --- |
| enriched | monocarboxylic acid metabolic process | 3.18E-017 |
| enriched | organic acid catabolic process&carboxylic acid catabolic process | 3.46E-017 |
| enriched | cellular amino acid catabolic process | 3.66E-017 |
| enriched | cellular amino acid metabolic process | 3.66E-017 |
| enriched | small molecule catabolic process | 5.60E-017 |
| enriched | alpha amino acid catabolic process | 9.35E-017 |
| enriched | alpha amino acid metabolic process | 2.87E-016 |
| enriched | small molecule biosynthetic process | 2.19E-014 |
| enriched | organic acid biosynthetic process&carboxylic acid biosynthetic process | 1.35E-013 |
| enriched | organonitrogen compound catabolic process | 1.18E-012 |
| enriched | regulation of body fluid levels | 1.18E-009 |
| enriched | wound healing | 1.18E-009 |
| enriched | anion transport | 3.32E-009 |
| enriched | liver development | 4.02E-009 |
| enriched | hepaticobiliary system development | 4.64E-009 |

(d) rat_muscle_DOWN

| **Type** | **Name** | **Adjusted p-value** |
| --- | --- | --- |
| enriched | blood vessel development | 2.90E-022 |
| enriched | regulation of cell migration | 2.90E-022 |
| enriched | vasculature development | 5.97E-022 |
| enriched | blood vessel morphogenesis | 1.56E-019 |
| enriched | regulation of cell morphogenesis | 1.75E-017 |
| enriched | cellular response to growth factor stimulus | 3.25E-017 |
| enriched | response to growth factor | 3.25E-017 |
| enriched | cell substrate adhesion | 5.25E-017 |
| enriched | tube development | 3.76E-016 |
| enriched | actin cytoskeleton organization | 8.75E-016 |
| enriched | extracellular matrix organization | 8.75E-016 |
| enriched | extracellular structure organization | 8.75E-016 |
| enriched | regulation of cell projection organization | 2.48E-015 |
| enriched | regulation of neuron differentiation | 4.67E-015 |
| enriched | regulation of cell morphogenesis involved in differentiation | 5.59E-015 |
| enriched | ossification | 1.14E-014 |
