## Supplementary material for "Age-associated different transcriptome profiling in zebrafish and rat: insight into diversity of vertebrate aging"

(a) rat_brain_UP

| **Type** | **Name** | **Adjusted p-value** |
| --- | --- | --- |
| enriched | response to drug | 1.62E-014 |
| enriched | response to peptide | 1.45E-012 |
| enriched | response to steroid hormone | 6.36E-012 |
| enriched | response to peptide hormone | 1.81E-011 |
| enriched | lipid localization | 1.04E-009 |
| enriched | cytokine production | 1.07E-009 |
| enriched | response to nutrient levels | 3.86E-009 |
| enriched | response to extracellular stimulus | 6.99E-009 |
| enriched | regulation of cytokine production | 1.14E-008 |
| enriched | response to corticosteroid | 1.81E-008 |
| enriched | lipid storage | 2.29E-008 |
| enriched | response to inorganic substance | 2.35E-008 |
| enriched | response to glucocorticoid | 8.37E-008 |
| enriched | response to oxygen levels | 2.81E-007 |
| enriched | blood vessel development | 3.29E-007 |

(b) rat_heart_UP

| **Type** | **Name** | **Adjusted p-value** |
| --- | --- | --- |
| enriched | regulation of cell projection organization | 6.73E-010 |
| enriched | regulation of secretion by cell | 1.13E-009 |
| enriched | regulation of system process | 1.13E-009 |
| enriched | wound healing | 1.16E-008 |
| enriched | developmental growth | 1.62E-008 |
| enriched | heart development | 1.76E-008 |
| enriched | regulation of cell migration | 1.76E-008 |
| enriched | regulation of neuron differentiation | 1.76E-008 |
| enriched | regulation of neuron projection development | 1.76E-008 |
| enriched | response to growth factor | 1.76E-008 |
| enriched | regulation of growth | 5.53E-008 |
| enriched | extracellular matrix organization | 8.11E-008 |
| enriched | extracellular structure organization | 8.11E-008 |
| enriched | cellular response to lipid | 8.62E-008 |
| enriched | gland development | 8.62E-008 |

(c) rat_liver_UP

| **Type** | **Name** | **Adjusted p-value** |
| --- | --- | --- |
| enriched | chromosome segregation | 1.54E-027 |
| enriched | mitotic cell cycle process | 1.32E-025 |
| enriched | mitotic nuclear division | 8.69E-024 |
| enriched | nuclear division | 8.69E-024 |
| enriched | organelle fission | 7.17E-023 |
| enriched | nuclear chromosome segregation | 3.49E-022 |
| enriched | mitotic sister chromatid segregation | 2.83E-019 |
| enriched | sister chromatid segregation | 6.35E-018 |
| enriched | cell division | 1.27E-016 |
| enriched | microtubule based process | 1.03E-015 |
| enriched | microtubule cytoskeleton organization | 1.18E-014 |
| enriched | regulation of chromosome segregation | 2.68E-014 |
| enriched | regulation of cell cycle process | 1.09E-012 |
| enriched | spindle organization | 1.13E-012 |
| enriched | regulation of cell division | 1.64E-012 |

(d) rat_muscle_UP

| **Type** | **Name** | **Adjusted p-value** |
| --- | --- | --- |
| enriched | monocarboxylic acid metabolic process | 6.67E-006 |
| enriched | fatty acid metabolic process | 1.20E-005 |
| enriched | fatty acid beta oxidation | 1.00E-004 |
| enriched | fatty acid catabolic process | 3.77E-004 |
| enriched | fatty acid oxidation | 4.60E-004 |
| enriched | lipid oxidation | 4.60E-004 |
| enriched | monocarboxylic acid catabolic process | 6.01E-004 |
| enriched | cellular lipid catabolic process | 1.31E-003 |
| enriched | lipid modification | 1.86E-003 |
| enriched | striated muscle contraction | 4.22E-003 |
| enriched | muscle system process | 1.81E-002 |
| enriched | lipid catabolic process | 2.03E-002 |
| enriched | organic acid catabolic process&carboxylic acid catabolic process | 2.03E-002 |
| enriched | small molecule catabolic process | 2.16E-002 |
