## Supplementary figures and images for "Age-associated different transcriptome profiling in zebrafish and rat: insight into diversity of vertebrate aging"

### Supplementary file 7

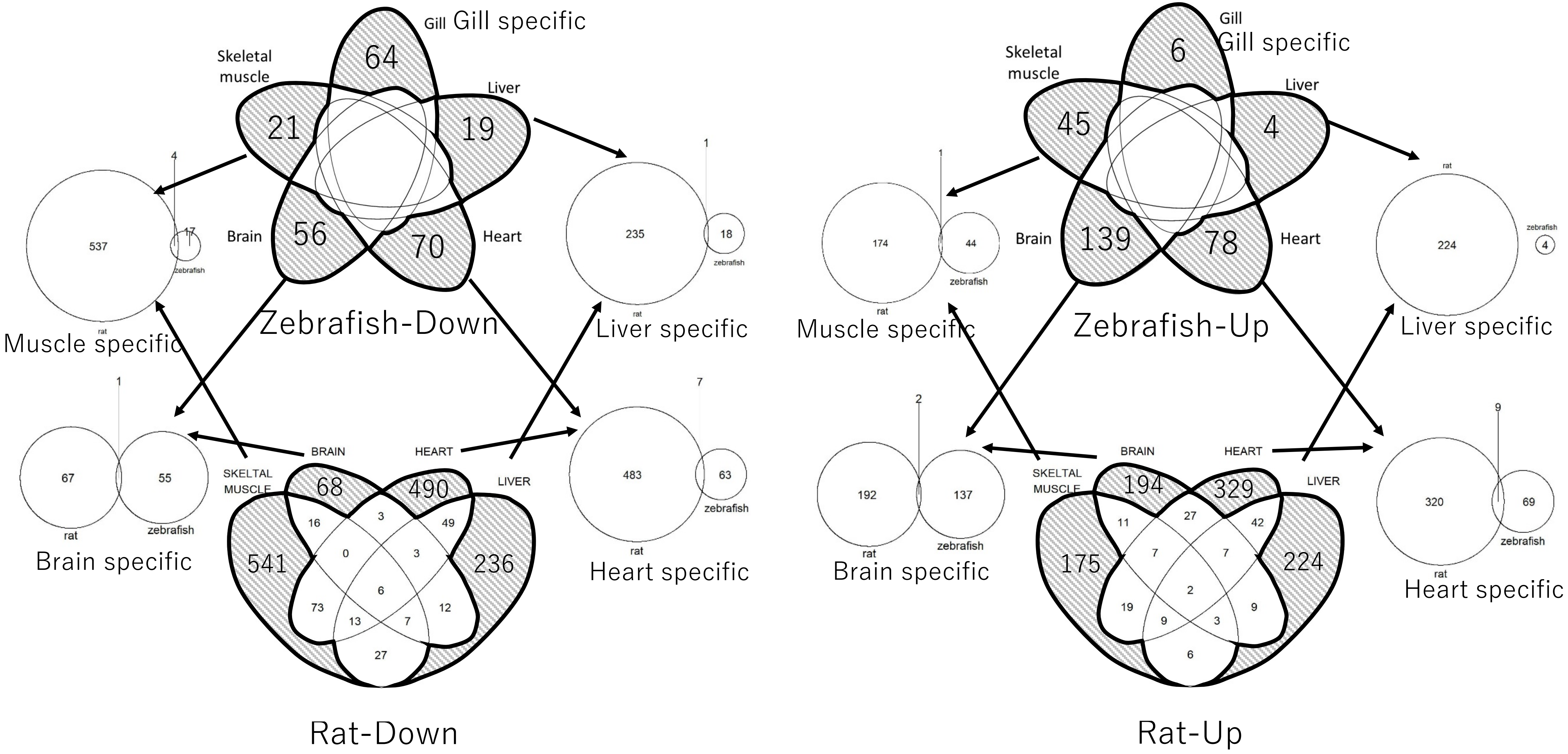

### Supplementary file 8

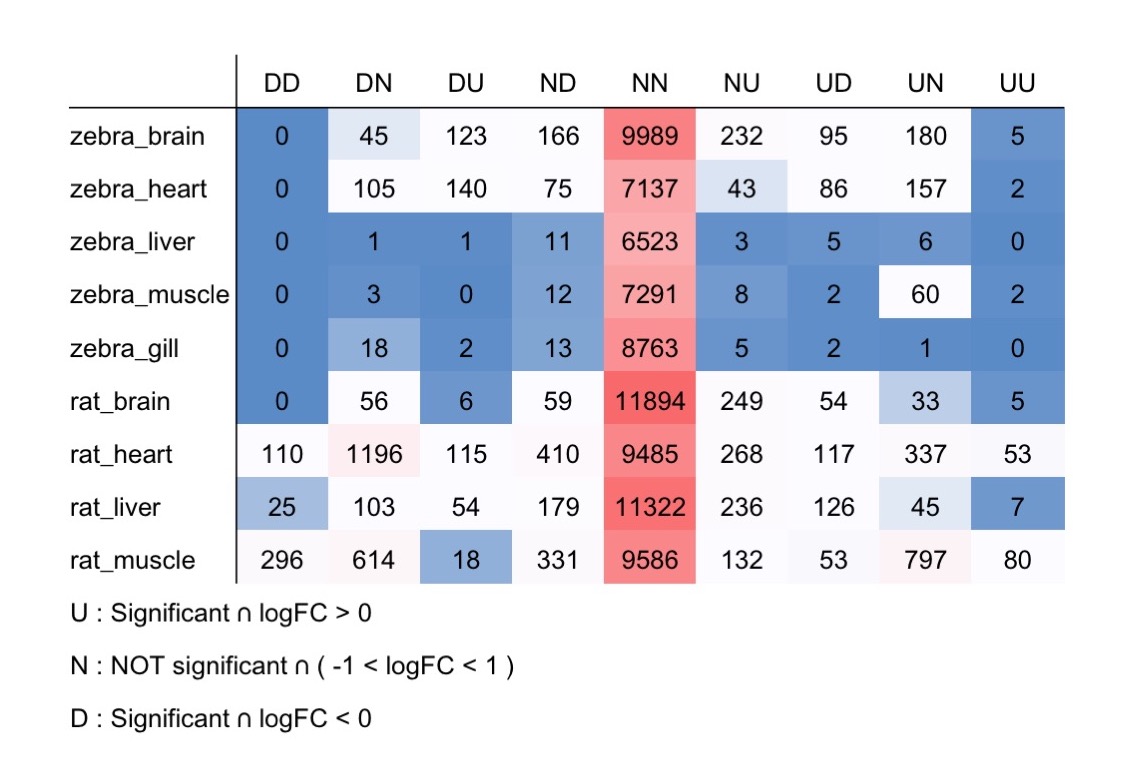

### Supplementary file 10

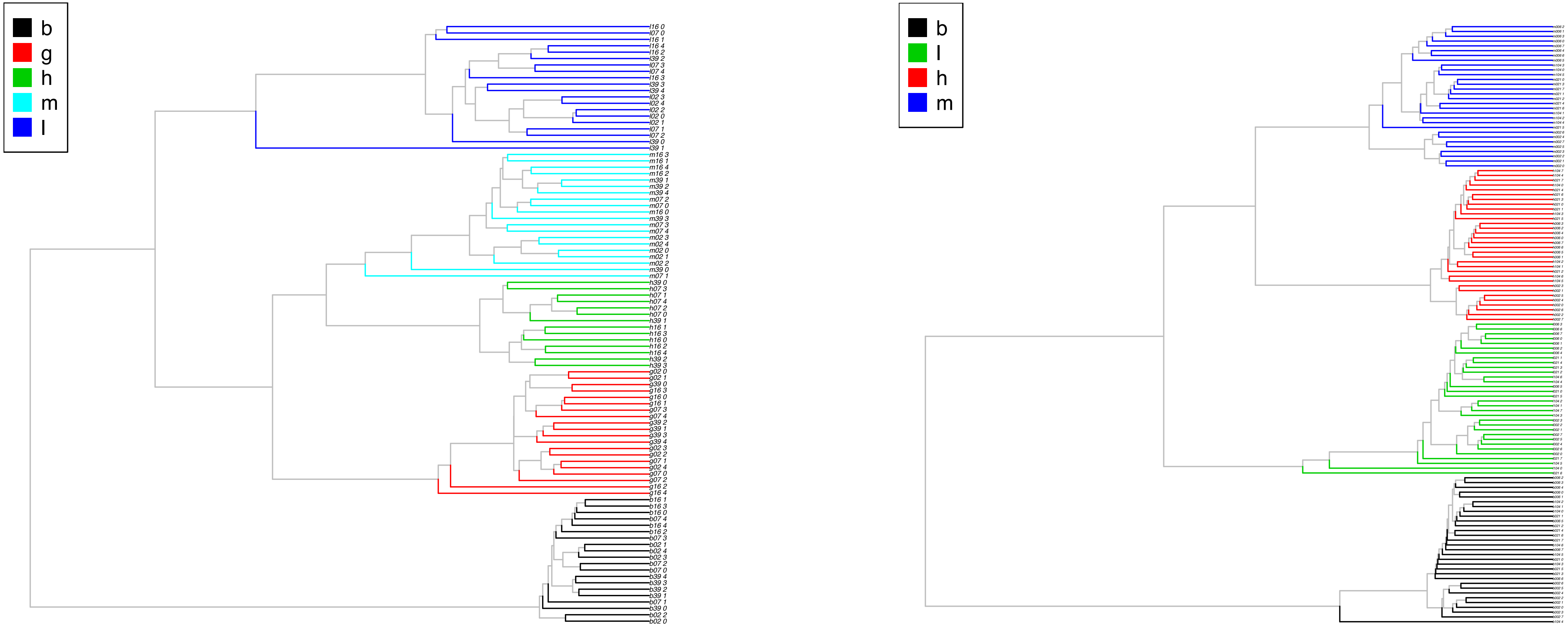


(a) Zebrafish

(b) Rat
