## Supplementary material for "Age-associated different transcriptome profiling in zebrafish and rat: insight into diversity of vertebrate aging"

| **Tissues** | **gene** | **sample_1** | **sample_2** | **fpkm_1** | **fpkm_2** | **log2_fold_change** | **q_value** |
| --- | --- | --- | --- | --- | --- | --- | --- |
| **Brain** | hif1al | b02 | b07 | 37.7893 | 10.385 | -1.86348 | 0.00204132 |
|  | nr1d2a | b02 | b07 | 63.4584 | 20.2966 | -1.64457 | 0.00204132 |
|  | hmgb3a | b02 | b39 | 629.699 | 213.047 | -1.56349 | 0.00204132 |
|  | hmgb3a | b02 | b07 | 629.699 | 239.548 | -1.39435 | 0.00204132 |
|  | klf13 | b02 | b07 | 35.5637 | 14.6073 | -1.28372 | 0.00204132 |
|  | csrnp1b | b02 | b07 | 35.1136 | 14.8692 | -1.23971 | 0.00204132 |
|  | pdk2a | b02 | b07 | 13.1941 | 5.95114 | -1.14866 | 0.00204132 |
|  | klf13 | b16 | b39 | 39.4604 | 18.6803 | -1.07889 | 0.00204132 |
|  | nr1d1 | b02 | b07 | 16.3326 | 7.7789 | -1.07012 | 0.00506054 |
|  | hmgb3a | b02 | b16 | 629.699 | 304.536 | -1.04805 | 0.00204132 |
|  | klf13 | b02 | b39 | 35.5637 | 18.6803 | -0.928892 | 0.00204132 |
|  | nr1d2a | b16 | b39 | 137.267 | 78.5585 | -0.805142 | 0.00204132 |
|  | csrnp1b | b16 | b39 | 49.7886 | 29.9604 | -0.73276 | 0.00204132 |
|  | pdk2a | b16 | b39 | 18.4573 | 12.0227 | -0.618429 | 0.0493388 |
|  | hmgb3a | b16 | b39 | 304.536 | 213.047 | -0.515443 | 0.0412412 |
|  | LOC566587 | b02 | b39 | 20.285 | 30.5406 | 0.590314 | 0.043526 |
|  | si:ch211-235e18.3 | b07 | b39 | 7.47557 | 11.5134 | 0.623062 | 0.048823 |
|  | slmo2 | b02 | b39 | 19.6702 | 32.1713 | 0.709764 | 0.0314399 |
|  | snrkb | b02 | b16 | 24.9048 | 40.7732 | 0.711196 | 0.00362646 |
|  | LOC566587 | b02 | b16 | 20.285 | 33.8836 | 0.740171 | 0.00362646 |
|  | LOC564054 | b02 | b16 | 10.5237 | 17.6255 | 0.744021 | 0.021629 |
|  | slmo2 | b16 | b39 | 18.9371 | 32.1713 | 0.764558 | 0.0223999 |
|  | snrkb | b07 | b16 | 23.5277 | 40.7732 | 0.793259 | 0.00636043 |
|  | crema | b07 | b39 | 10.0733 | 17.7955 | 0.820985 | 0.00869823 |
|  | slmo2 | b07 | b39 | 17.1414 | 32.1713 | 0.908289 | 0.00362646 |
|  | crema | b02 | b39 | 9.44241 | 17.7955 | 0.91429 | 0.00204132 |
|  | si:ch211-235e18.3 | b02 | b39 | 5.94453 | 11.5134 | 0.953682 | 0.00204132 |
|  | csrnp1b | b07 | b39 | 14.8692 | 29.9604 | 1.01073 | 0.00204132 |
|  | pdk2a | b07 | b39 | 5.95114 | 12.0227 | 1.01453 | 0.00204132 |
|  | crema | b07 | b16 | 10.0733 | 20.3509 | 1.01456 | 0.00204132 |
|  | ulk2 | b07 | b16 | 9.01704 | 18.5129 | 1.03781 | 0.00204132 |
|  | si:ch211-235e18.3 | b07 | b16 | 7.47557 | 15.7652 | 1.07649 | 0.00204132 |
|  | si:ch1073-281m9.1 | b02 | b39 | 7.20986 | 15.4092 | 1.09575 | 0.0177734 |
|  | crema | b02 | b16 | 9.44241 | 20.3509 | 1.10786 | 0.00204132 |
|  | nr1d2a | b02 | b16 | 63.4584 | 137.267 | 1.1131 | 0.00204132 |
|  | LOC566587 | b07 | b39 | 13.9427 | 30.5406 | 1.13122 | 0.00204132 |
|  | LOC564054 | b07 | b39 | 5.95822 | 13.8745 | 1.21948 | 0.00204132 |
|  | LOC566587 | b07 | b16 | 13.9427 | 33.8836 | 1.28107 | 0.00204132 |
|  | si:ch211-235e18.3 | b02 | b16 | 5.94453 | 15.7652 | 1.40711 | 0.00204132 |
|  | klf13 | b07 | b16 | 14.6073 | 39.4604 | 1.43372 | 0.00204132 |
|  | hif1al | b07 | b39 | 10.385 | 29.2568 | 1.49428 | 0.00204132 |
|  | LOC564054 | b07 | b16 | 5.95822 | 17.6255 | 1.56471 | 0.00204132 |
|  | pdk2a | b07 | b16 | 5.95114 | 18.4573 | 1.63296 | 0.00204132 |
|  | nr1d1 | b02 | b16 | 16.3326 | 54.4783 | 1.73793 | 0.00204132 |
|  | csrnp1b | b07 | b16 | 14.8692 | 49.7886 | 1.74349 | 0.00204132 |
|  | hif1al | b07 | b16 | 10.385 | 38.8428 | 1.90315 | 0.00204132 |
|  | nr1d2a | b07 | b39 | 20.2966 | 78.5585 | 1.95253 | 0.00204132 |
|  | nr1d1 | b02 | b39 | 16.3326 | 64.3612 | 1.97844 | 0.00204132 |
|  | fos | b02 | b07 | 5.65843 | 25.0391 | 2.14571 | 0.00204132 |
|  | fos | b02 | b39 | 5.65843 | 28.8997 | 2.35258 | 0.00204132 |
|  | fos | b02 | b16 | 5.65843 | 32.7026 | 2.53093 | 0.00204132 |
|  | nr1d2a | b07 | b16 | 20.2966 | 137.267 | 2.75767 | 0.00204132 |
|  | nr1d1 | b07 | b16 | 7.7789 | 54.4783 | 2.80804 | 0.00204132 |
|  | pdk2b | b07 | b16 | 1.01042 | 7.55262 | 2.90203 | 0.00204132 |
|  | pdk2b | b07 | b39 | 1.01042 | 7.87189 | 2.96176 | 0.00204132 |
|  | nr1d1 | b07 | b39 | 7.7789 | 64.3612 | 3.04855 | 0.00204132 |
| **Heart** | pdk2a | h16 | h39 | 175.584 | 47.6919 | -1.88035 | 0.00417475 |
|  | si:ch211-235e18.3 | h16 | h39 | 31.0838 | 13.0984 | -1.24678 | 0.0471188 |
|  | hmgb3a | h16 | h39 | 41.533 | 19.0254 | -1.12633 | 0.0272986 |
|  | LOC564054 | h07 | h16 | 10.1832 | 24.6667 | 1.27638 | 0.0303784 |
|  | slmo2 | h07 | h16 | 59.2739 | 179.526 | 1.59872 | 0.00519275 |
|  | crema | h07 | h39 | 7.53261 | 23.5649 | 1.64542 | 0.023094 |
|  | ulk2 | h07 | h39 | 2.2118 | 6.95761 | 1.65337 | 0.00519275 |
|  | snrkb | h07 | h39 | 8.55654 | 27.9737 | 1.70897 | 0.0253635 |
|  | crema | h07 | h16 | 7.53261 | 24.7843 | 1.71821 | 0.012957 |
|  | pdk2b | h07 | h16 | 41.769 | 142.769 | 1.77318 | 0.0211518 |
|  | pdk2a | h07 | h16 | 46.4564 | 175.584 | 1.91821 | 0.00175208 |
|  | klf13 | h07 | h39 | 2.03789 | 7.97645 | 1.96867 | 0.0123385 |
|  | pdk2b | h07 | h39 | 41.769 | 169.924 | 2.02438 | 0.00798355 |
|  | hif1al | h07 | h39 | 8.7201 | 36.5921 | 2.06912 | 0.00175208 |
|  | si:ch1073-281m9.1 | h07 | h39 | 110.005 | 471.044 | 2.09829 | 0.0123385 |
|  | slmo2 | h07 | h39 | 59.2739 | 273.384 | 2.20546 | 0.00175208 |
|  | fos | h16 | h39 | 14.9062 | 72.3957 | 2.27999 | 0.00175208 |
|  | ulk2 | h07 | h16 | 2.2118 | 12.33 | 2.47888 | 0.00175208 |
|  | nr1d1 | h07 | h16 | 3.3309 | 19.4953 | 2.54914 | 0.00175208 |
|  | si:ch1073-281m9.1 | h16 | h39 | 76.2942 | 471.044 | 2.62622 | 0.00304614 |
|  | hif1al | h07 | h16 | 8.7201 | 61.4973 | 2.81811 | 0.00175208 |
|  | klf13 | h07 | h16 | 2.03789 | 14.4941 | 2.83032 | 0.00175208 |
|  | csrnp1b | h07 | h39 | 6.76973 | 49.3229 | 2.86509 | 0.00175208 |
|  | LOC566587 | h07 | h39 | 3.03895 | 22.5919 | 2.89416 | 0.00175208 |
|  | snrkb | h07 | h16 | 8.55654 | 65.349 | 2.93307 | 0.00175208 |
|  | nr1d2a | h07 | h39 | 4.95923 | 46.1528 | 3.21823 | 0.00175208 |
|  | LOC566587 | h07 | h16 | 3.03895 | 33.3483 | 3.45597 | 0.00175208 |
|  | csrnp1b | h07 | h16 | 6.76973 | 74.6757 | 3.46347 | 0.00175208 |
|  | fos | h07 | h39 | 5.60336 | 72.3957 | 3.69154 | 0.00175208 |
|  | nr1d1 | h07 | h39 | 3.3309 | 44.9769 | 3.7552 | 0.00175208 |
|  | nr1d2a | h07 | h16 | 4.95923 | 72.1624 | 3.86306 | 0.00175208 |
| **Liver** | hmgb3a | l02 | l07 | 8.10787 | 1.37502 | -2.55987 | 0.0343286 |
|  | nr1d1 | l07 | l16 | 5.28559 | 26.3746 | 2.31901 | 0.0457057 |
|  | csrnp1b | l07 | l16 | 2.75304 | 15.9449 | 2.534 | 0.0343286 |
|  | si:ch1073-281m9.1 | l02 | l07 | 2.37994 | 15.7912 | 2.73013 | 0.0372533 |
|  | slmo2 | l07 | l16 | 15.8092 | 117.269 | 2.89098 | 0.0343286 |
|  | csrnp1b | l02 | l16 | 1.98091 | 15.9449 | 3.00886 | 0.0150697 |
|  | LOC564054 | l07 | l16 | 0.999058 | 8.71313 | 3.12455 | 0.00888127 |
|  | fos | l02 | l16 | 9.48711 | 88.0693 | 3.2146 | 0.0150697 |
|  | fos | l07 | l16 | 9.40857 | 88.0693 | 3.22659 | 0.0150697 |
|  | LOC566587 | l07 | l16 | 6.6533 | 66.253 | 3.31584 | 0.0150697 |
|  | si:ch1073-281m9.1 | l02 | l39 | 2.37994 | 26.2787 | 3.4649 | 0.00888127 |
|  | klf13 | l07 | l16 | 4.52957 | 61.428 | 3.76145 | 0.0150697 |
|  | crema | l02 | l16 | 3.53388 | 54.0853 | 3.93591 | 0.00888127 |
|  | hif1al | l07 | l39 | 2.17032 | 34.41 | 3.98685 | 0.00888127 |
|  | pdk2a | l02 | l16 | 4.43019 | 78.3313 | 4.14415 | 0.00888127 |
|  | nr1d2a | l07 | l16 | 3.546 | 63.565 | 4.16397 | 0.0278497 |
|  | ulk2 | l07 | l16 | 1.93809 | 38.444 | 4.31005 | 0.00888127 |
|  | si:ch211-235e18.3 | l02 | l16 | 1.29696 | 29.703 | 4.5174 | 0.00888127 |
|  | pdk2b | l07 | l16 | 6.20329 | 182.07 | 4.87531 | 0.0372533 |
|  | snrkb | l07 | l16 | 1.37763 | 47.5333 | 5.10868 | 0.00888127 |
| **Muscle** | hmgb3a | m02 | m39 | 27.6871 | 2.41291 | -3.52037 | 0.00533534 |
|  | fos | m02 | m07 | 34.8488 | 3.07734 | -3.50135 | 0.00881964 |
|  | hif1al | m02 | m07 | 24.7479 | 5.74447 | -2.10706 | 0.0410281 |
|  | hmgb3a | m16 | m39 | 10.1069 | 2.41291 | -2.06649 | 0.0453296 |
|  | slmo2 | m02 | m07 | 135.833 | 35.103 | -1.95217 | 0.0405262 |
|  | klf13 | m02 | m07 | 4.68521 | 1.55396 | -1.59216 | 0.0349222 |
|  | crema | m02 | m39 | 4.09367 | 11.4447 | 1.48321 | 0.0449324 |
|  | crema | m07 | m16 | 1.87112 | 6.28319 | 1.7476 | 0.0421944 |
|  | LOC566587 | m07 | m16 | 2.4719 | 9.44699 | 1.93423 | 0.0239486 |
|  | LOC566587 | m07 | m39 | 2.4719 | 9.96984 | 2.01195 | 0.0156865 |
|  | slmo2 | m07 | m39 | 35.103 | 148.515 | 2.08095 | 0.0174025 |
|  | nr1d1 | m02 | m39 | 4.3258 | 19.8136 | 2.19545 | 0.0388622 |
|  | csrnp1b | m07 | m39 | 2.96351 | 14.0926 | 2.24956 | 0.00533534 |
|  | klf13 | m07 | m16 | 1.55396 | 7.42855 | 2.25713 | 0.0213414 |
|  | csrnp1b | m07 | m16 | 2.96351 | 14.3644 | 2.27712 | 0.00533534 |
|  | ulk2 | m02 | m16 | 1.8134 | 9.49404 | 2.38833 | 0.00533534 |
|  | si:ch211-235e18.3 | m07 | m39 | 6.83552 | 39.0204 | 2.5131 | 0.0138016 |
|  | ulk2 | m02 | m39 | 1.8134 | 10.5148 | 2.53566 | 0.0342146 |
|  | fos | m07 | m16 | 3.07734 | 17.9325 | 2.54282 | 0.0116017 |
|  | crema | m07 | m39 | 1.87112 | 11.4447 | 2.6127 | 0.00533534 |
|  | pdk2a | m02 | m39 | 2.67909 | 17.3525 | 2.69533 | 0.00533534 |
|  | fos | m07 | m39 | 3.07734 | 20.4279 | 2.73079 | 0.00533534 |
|  | si:ch1073-281m9.1 | m02 | m39 | 8.02844 | 58.8295 | 2.87335 | 0.0349222 |
|  | hif1al | m07 | m39 | 5.74447 | 44.8833 | 2.96593 | 0.0325192 |
|  | snrkb | m07 | m16 | 1.907 | 15.0074 | 2.9763 | 0.0138016 |
|  | si:ch211-235e18.3 | m07 | m16 | 6.83552 | 54.0787 | 2.98394 | 0.0342146 |
|  | nr1d2a | m07 | m39 | 3.3532 | 27.8179 | 3.0524 | 0.0187025 |
|  | pdk2a | m02 | m16 | 2.67909 | 23.9243 | 3.15866 | 0.00533534 |
|  | slmo2 | m07 | m16 | 35.103 | 322.041 | 3.19758 | 0.00533534 |
|  | si:ch1073-281m9.1 | m07 | m16 | 2.45601 | 23.1627 | 3.23741 | 0.0116017 |
|  | pdk2b | m02 | m39 | 19.1174 | 181.39 | 3.24614 | 0.00533534 |
|  | nr1d1 | m02 | m16 | 4.3258 | 41.4588 | 3.26064 | 0.00533534 |
|  | ulk2 | m07 | m16 | 0.901769 | 9.49404 | 3.39619 | 0.00533534 |
|  | ulk2 | m07 | m39 | 0.901769 | 10.5148 | 3.54352 | 0.00533534 |
|  | LOC564054 | m07 | m16 | 0.615569 | 7.1904 | 3.54608 | 0.0453296 |
|  | pdk2b | m02 | m16 | 19.1174 | 259.482 | 3.76268 | 0.00533534 |
|  | hif1al | m07 | m16 | 5.74447 | 99.8005 | 4.1188 | 0.041509 |
|  | nr1d2a | m07 | m16 | 3.3532 | 61.8209 | 4.20448 | 0.00533534 |
|  | pdk2b | m07 | m39 | 9.28608 | 181.39 | 4.28788 | 0.00533534 |
|  | si:ch1073-281m9.1 | m07 | m39 | 2.45601 | 58.8295 | 4.58215 | 0.00533534 |
|  | pdk2b | m07 | m16 | 9.28608 | 259.482 | 4.80442 | 0.00533534 |
| **Gill** | hmgb3a | g16 | g39 | 70.8538 | 20.919 | -1.76003 | 0.0366772 |
|  | LOC566587 | g16 | g39 | 11.2948 | 47.5359 | 2.07336 | 0.00814895 |
